# A disease dynamics atlas forecasts patient states and maps molecular programs in ALS

**DOI:** 10.64898/2026.02.01.703078

**Authors:** Zizhang Li, Chong Gao, Jiaming Kong, Yuting Fu, Shan Wen, Guangpeng Li, Ye Cao, Yunlin Fu, Hao Zhang, Shikai Jia, Xiaojing Liu, Jiangli Yang, Lei Cai, Feng Yan, Xiaodong Liu, Luyi Tian

**Author notes:** These authors contributed equally to this work (Zizhang Li, Chong Gao, Jiaming Kong, Yuting Fu).

## Abstract

ALS progression is multidimensional, yet fragmented records and scalar outcomes obscure how patients move through disease states and how those states relate to molecular variation. MEDSTREM converts patient-held medical-record images into standardised longitudinal data, enabling bottom-up cohort construction. Using MEDSTREM-structured records from more than 8,000 AskHelpU participants together with PRO-ACT and Answer ALS, we developed DynaALS, the ALS Disease Dynamics Atlas. DynaALS represents ALS as a dynamic patient-state manifold that captures distinct directions of deterioration and patient movement between them over time. DynaALS retrieved population-referenced future states and decoded them into multidimensional clinical profiles without requiring patient-specific longitudinal histories. Motor-neuron RNA and chromatin profiles linked DynaALS states to developmental and regulatory programs, while neuromuscular-organoid single-cell multi-omics converged on a neural-developmental Netrin-DCC signalling axis across interacting cell types. By coupling MEDSTREM-enabled data construction to dynamic state modelling, DynaALS establishes a transferable patient-state engine for predictive and biologically interpretable disease models.

## Main

Amyotrophic lateral sclerosis (ALS) is a fatal neurodegenerative disease marked by pronounced heterogeneity in site of onset, rate of functional decline and survival, complicating prognosis, endpoint selection and trial design^1,2^. The ALS Functional Rating Scale-Revised (ALSFRS-R) remains the dominant longitudinal outcome^3,4^. Its total score and slope, however, compress distinct clinical domains into one scalar and are sensitive to sparse, irregular follow-up^5^. King’s and Milano-Torino staging provide interpretable milestones based on regional involvement or loss of independence^6,7^. They emphasise different disease phases but retain limited continuous information within stages^8^. ALS therefore needs a representation that preserves both how far disease has progressed and the clinical route by which deterioration unfolds.

Resources and modern models have addressed parts of this problem. PRO-ACT standardised longitudinal trial data^9^, whereas Answer ALS linked clinical phenotyping to multi-omics from induced pluripotent stem cell-derived motor neurons^10^. Trajectory, latent-variable, manifold-learning, Bayesian and neural-network models have identified progression groups, simulated impairment or predicted selected outcomes^11–17^. Molecular studies have also described RNA and chromatin heterogeneity in patient-derived motor neurons^10,18^. These approaches solve important but separate tasks, including stratification, trajectory clustering, outcome prediction and molecular association. What remains missing is a reusable patient-time geometry that unifies these tasks without collapsing ALS into one outcome. Such a geometry should separate global burden from domain composition, represent longitudinal movement and support several clinical and molecular readouts.

A second limitation lies in how ALS cohorts are constructed. Most ALS models start with records already structured by trials or specialist centres, while real-world histories remain dispersed across hospitals and retained by patients as reports, scans or photographs. AskHelpU offers a different starting point. Through mobile and web interfaces, individuals upload documents from their own care histories and contribute dated ALSFRS-R assessments. More than 8,000 individuals have contributed. Few ALS resources begin with patient-held document images at this scale, making AskHelpU a distinct bottom-up complement to trial and specialist-centre cohorts. Yet collecting documents is not equivalent to constructing analysis-ready longitudinal data: dates, terminology, units and provenance must be reconciled into auditable records that can be harmonised across cohorts. A useful ALS atlas therefore requires both a patient-mediated data pathway and a reusable state-space model.

We addressed these coupled needs with MEDSTREM and DynaALS. MEDSTREM is a patient-mediated, locally deployable system that uses large language models to transform patient-sourced medical-record images into standardised longitudinal health records within authorised infrastructure. Its companion application supports upload, review, editing, timeline visualisation and export, returning a usable health record to the individual while enabling bottom-up cohort construction. We harmonised AskHelpU records with PRO-ACT and Answer ALS to create DynaALS, the ALS Disease Dynamics Atlas, a shared disease-state manifold in which serial observations trace patient movement through ALS progression. Unlike a patient-level summary or fixed subtype, this patient-time representation preserves within-patient change and allows clinical composition to be remapped as disease progresses. Within one coordinate system, DynaALS separates global burden from functional, respiratory, nutritional and mixed clinical directions and captures their temporal reorganisation. It retrieves population-referenced future states and decodes forecast manifold positions into multidimensional clinical profiles without requiring a patient’s longitudinal history. As new observations accrue, they update patient positions and forecasts without rebuilding the atlas. The same coordinates link clinical directions to RNA and chromatin programs in induced motor neurons. Single-cell multi-omics of neuromuscular organoids then resolves these programs across interacting cell types. This cross-scale mapping converged on a neural-developmental Netrin-DCC signalling axis and prioritised it for perturbational testing. The central advance is to couple locally controlled data construction to a shared atlas in which ALS heterogeneity becomes dynamic, queryable and biologically interpretable. Together, MEDSTREM and DynaALS establish a new research paradigm for constructing continuously updated virtual-patient models: longitudinal records locate an individual in disease space, population dynamics forecast future states, and each new observation refreshes the representation. Developed in ALS, this framework advances clinically grounded digital twins and offers a transferable strategy for other diseases.

## Results

We first used MEDSTREM to transform patient-uploaded AskHelpU records into standardised longitudinal data (Fig. 1a). We then integrated these data with Answer ALS and PRO-ACT to construct DynaALS, the ALS Disease Dynamics Atlas, a shared patient-state manifold that resolves clinical heterogeneity and supports population-referenced future-state retrieval and multidimensional clinical decoding. Finally, we linked DynaALS-defined clinical states to molecular programs in induced motor neurons (iMNs) and neuromuscular organoids (NMOs).

**Fig. 1.**
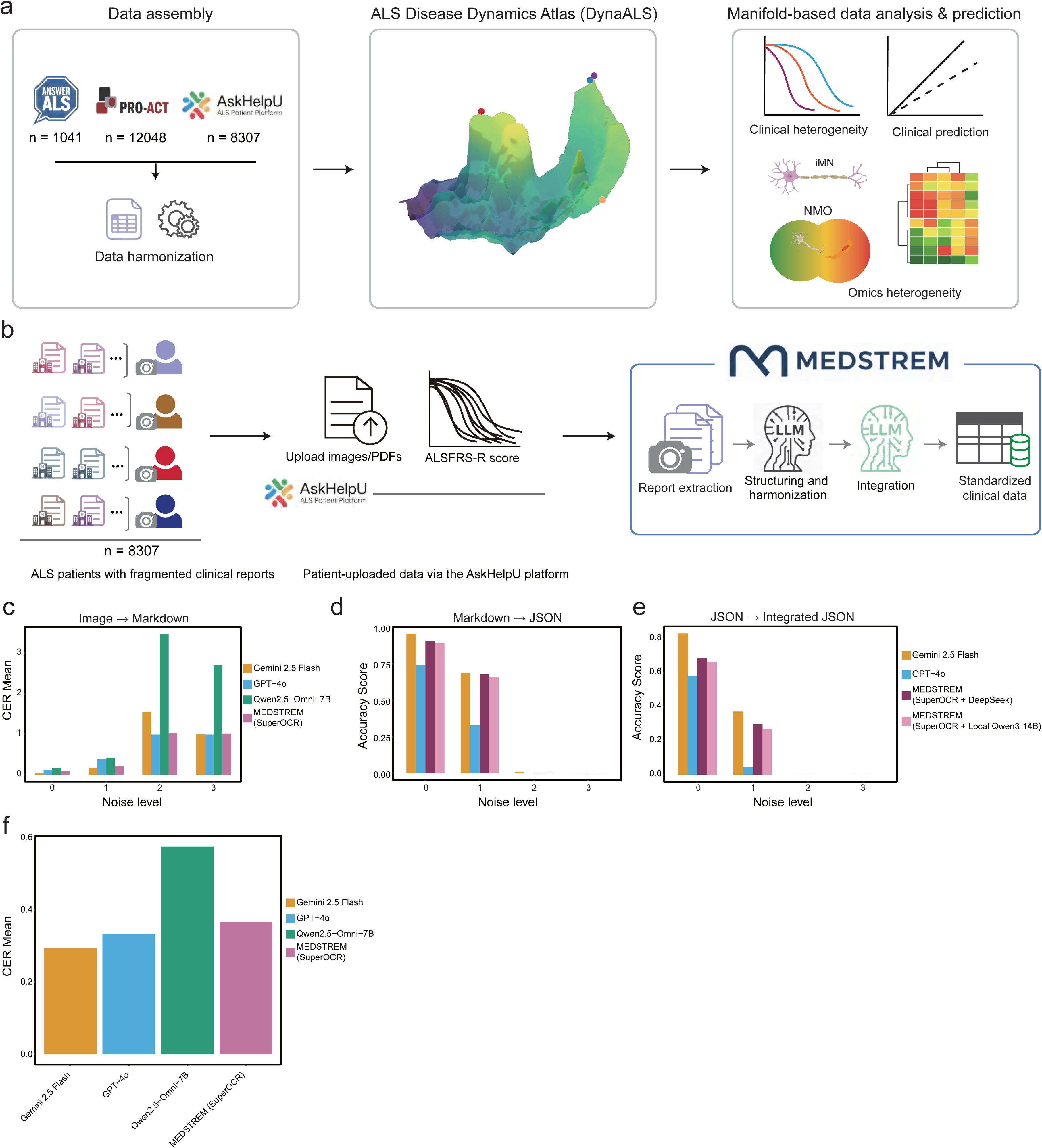
MEDSTREM enables patient-mediated construction of longitudinal ALS records. a. Study overview linking patient-mediated AskHelpU intake with Answer ALS and PRO-ACT through DynaALS, the ALS Disease Dynamics Atlas, and patient-derived molecular models. b. AskHelpU-to-structured-record workflow. AskHelpU combined linked document-upload and structured-assessment channels. c. Mean character error rate (CER; lower is better) across four synthetic noise levels for Gemini 2.5 Flash, GPT-4o, Qwen2.5-Omni-7B and MEDSTREM SuperOCR. Each model-by-noise group comprised 1,980-2,000 images. d. Mean flattened key-path field accuracy (higher is better) for document-JSON extraction using Gemini 2.5 Flash, GPT-4o, MEDSTREM with DeepSeek V3 or local Qwen3-14B. e. Mean patient-level key-path accuracy after date-aware integration for four configurations in 20 synthetic patients per noise level. f. Mean CER on manually deidentified real clinic-note or vital-sign, electrocardiography, electromyography and laboratory records.

### An agentic framework enables privacy-preserving, large-scale clinical data ingestion

AskHelpU enabled a patient-mediated model of ALS cohort construction (Fig. 1a,b). More than 8,000 participants used WeChat and web interfaces to aggregate patient-held files and photographs from dispersed care encounters and contribute dated ALSFRS-R assessments (Extended Data Fig. 1a–c). In the source-coverage snapshot, participants contributed a mean of 19.5 documents spanning 15.4 examination types, including diagnostic, imaging, electrophysiological, laboratory, medication and pulmonary records. This routine-care breadth complemented trial-focused PRO-ACT (mean, 9.5 types) and the denser protocol-defined phenotyping in Answer ALS (mean, 26.1 types; Extended Data Fig. 4). AskHelpU therefore established a continuously expanding, patient-contributed source of real-world ALS evidence at cohort scale.

MEDSTREM transformed these heterogeneous contributions into reviewable longitudinal histories (Fig. 1b and Extended Data Fig. 2). SuperOCR combined orientation correction and conventional OCR guidance with a locally fine-tuned Qwen2.5-Omni-7B model trained on synthetic clinical reports degraded to reproduce rotation, blur, geometric distortion and background variation. It reconstructed headings, tables, dates, numerical values and units as structured Markdown. Downstream language-model agents converted each document into JSON, normalised terminology and units, reconciled observations by examination date and stored records with provenance and quality flags in MongoDB. A fully local SuperOCR–Qwen3-14B configuration kept the complete image-to-record chain within authorised infrastructure. The same core supported cohort-scale processing and an individual-facing application for upload, review, correction, timeline visualisation and export (Extended Data Fig. 13a). MEDSTREM thus combined decentralised contribution with auditable local computation, returning editable records to individuals while constructing a common ALS research resource.

We benchmarked the complete record-construction chain because image processing and document layout can jointly affect information recovery^19^. The synthetic benchmark comprised 100 patients, five document classes and four image-corruption levels, with a separate manually deidentified real-record evaluation (Fig. 1c–f and Extended Data Fig. 3). Consistent with task-specific adaptation of compact clinical models^20^, fine-tuning markedly improved SuperOCR over the underlying Qwen2.5-Omni-7B model. Under clean and light corruption, mean character error rate fell from 0.1618 to 0.0986 and from 0.4130 to 0.2097, relative reductions of 39% and 49%, respectively. SuperOCR outperformed GPT-4o under both conditions and approached Gemini 2.5 Flash (Fig. 1c). Performance declined for all systems at the two highest corruption levels. On real records, SuperOCR retained an advantage over the untuned baseline (0.3652 versus 0.5745) and approached GPT-4o (0.3337) and Gemini 2.5 Flash (0.2931; Fig. 1f).

Competitive local performance extended to structured record construction. Under clean and light corruption, local Qwen3-14B achieved document-level key-path accuracies of 0.8912 and 0.6596. These values exceeded GPT-4o (0.7423 and 0.3331) and closely matched DeepSeek V3 (0.9056 and 0.6785; Fig. 1d and Extended Data Fig. 3b). After longitudinal integration, local Qwen3-14B again exceeded GPT-4o (0.6423 and 0.2564 versus 0.5796 and 0.0428) and remained within 0.03 of DeepSeek V3 (0.6656 and 0.2836; Fig. 1e). The fully local SuperOCR-Qwen3 configuration therefore outperformed GPT-4o across all three stages under clean and light corruption. MEDSTREM extended prior local clinical information extraction to a complete image-to-longitudinal-record workflow, achieving competitive accuracy with workstation-scale models while protected records remained inside authorised infrastructure^21^.

### DynaALS creates a reusable geometry of multidimensional disease states

We harmonised MEDSTREM-structured AskHelpU records with existing longitudinal observations from Answer ALS and PRO-ACT. To establish a common global progression anchor, we derived the ALS Integrated Deterioration Index (AIDI), a participant-level measure integrating full-follow-up functional, respiratory and nutritional deterioration. In same-patient comparisons, AIDI correlated more strongly than ALSFRS-R decline and two stage-like summaries with FVC decline, SVC decline and weight loss (rho = 0.711, 0.755 and 0.325, respectively), while its association with grip-strength decline was comparable to that of ALSFRS-R decline (Extended Data Fig. 5b and Supplementary Table S4). These results support integrating complementary clinical domains into a common measure of overall ALS progression, while retaining domain-specific information for the disease-state representation that follows^22^. We next benchmarked AIDI against mixture-of-Gaussian-process (MoGP) modelling, an established ALSFRS-R trajectory-stratification algorithm that identifies nonlinear progression patterns from sparse longitudinal measurements^11^. Across all three resources, AIDI increased with the deterioration ordering of trajectories assigned to the published MoGP reference clusters (rho = 0.758-0.952; Extended Data Fig. 5c).

We then represented each longitudinal history as a series of reference-day patient-time states (Fig. 2a and Extended Data Fig. 4). State construction integrated functional, respiratory, nutritional, motor, neurological, vital-sign and laboratory observations, together with their longitudinal changes, recency, coverage and observation context. A partitioned encoder jointly modelled shared clinical status and patterns of clinical observation and data availability, using AIDI and complementary global and domain-specific deterioration targets to supervise the clinical representation. Subsequent metric refinement organised clinically similar states while preserving progression ordering across resources.

**Fig. 2.**
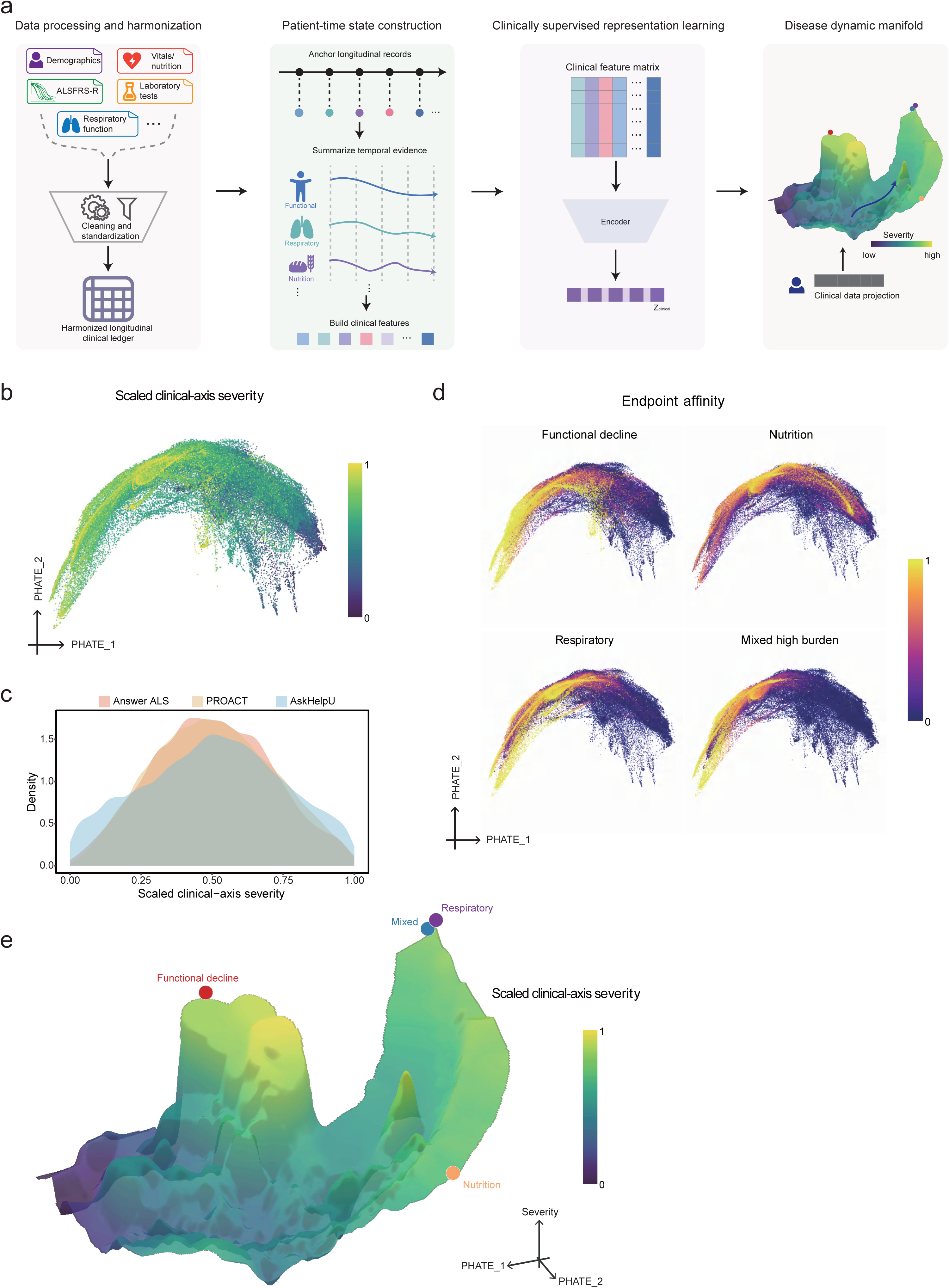
DynaALS creates a shared geometry of multidimensional disease states. a. Construction of DynaALS as the state-space engine of a clinical digital twin. MEDSTREM-structured AskHelpU records were harmonised with longitudinal observations from Answer ALS and PRO-ACT and summarised at patient-time landmarks. The state representation combined 42 clinical concepts with longitudinal change, coverage and observation-context features. A partitioned encoder jointly modelled shared clinical status and patterns of clinical observation and data availability under AIDI and complementary global and domain-specific supervision, followed by patient-grouped metric refinement. b. PHATE visualisation of 158,300 patient-time states, coloured by scaled domain severity from functional, respiratory and nutritional measurements; this state-level colouring is distinct from the participant-level ALS Integrated Deterioration Index (AIDI). c. Kernel-density distributions of scaled severity for Answer ALS (1,016 participants; 7,192 states), PRO-ACT (12,048 participants; 88,462 states) and AskHelpU (8,307 participants; 62,646 states). The Answer ALS subset comprised 848 participants with ALS, 88 healthy controls, 12 asymptomatic ALS gene carriers and 68 participants with non-ALS motor neuron disease. d. Graph-smoothed enrichment of functional, respiratory, nutritional and mixed high-burden reference states using Gaussian-distance-weighted neighbours on the 50-nearest-neighbour graph. e. Three-dimensional PHATE display of the clinical-burden surface and domain-specific high-burden affinity peaks. Neighbourhoods and affinity fields were computed in the full DynaALS representation and visualised using PHATE.

The resulting DynaALS organised 158,300 longitudinal clinical states from 21,371 participants into a shared geometry of ALS progression. Answer ALS contributed 7,192 states from 1,016 participants, PRO-ACT contributed 88,462 states from 12,048 participants and AskHelpU contributed 62,646 states from 8,307 participants (Fig. 2b,c). Longitudinal progression appeared as movement of patient states through this geometry, which we visualised using PHATE. Earlier manifold analyses resolved baseline prognostic zones or patient-level clinical subgroups^14,23^. DynaALS advances this strategy by organising repeated patient-time states and their movement through one shared geometry.

Functional, respiratory and nutritional severity could be reconstructed in held-out states using reference states from the other cohorts, with mean correlations of r = 0.910 in Answer ALS, 0.879 in PRO-ACT and 0.838 in AskHelpU, matching or exceeding within-cohort performance (Extended Data Fig. 6a). Graph-smoothed affinities delineated functional, respiratory, nutritional and mixed high-burden regions (Fig. 2d,e), and the three domain-specific affinities correlated with their corresponding clinical endpoints at rho = 0.842, 0.846 and 0.808 (Extended Data Fig. 6b,c). DynaALS expands the global progression axis captured by AIDI into a multidimensional geometry that represents both how far ALS has progressed and the clinical direction in which deterioration is unfolding.

### DynaALS reveals clinically distinct, dynamic directions of deterioration

Existing ALS studies have identified patient-level clinical subtypes and longitudinal progression clusters^11,12,23^. DynaALS added a state-level view by assigning direction to each patient-time state within a continuous disease geometry. Global severity described how far deterioration had advanced, whereas direction described its functional, respiratory, nutritional or mixed composition (Fig. 3a,b). Low-confidence boundary states remained uncertain (Methods; Supplementary Table S5a).

**Fig. 3.**
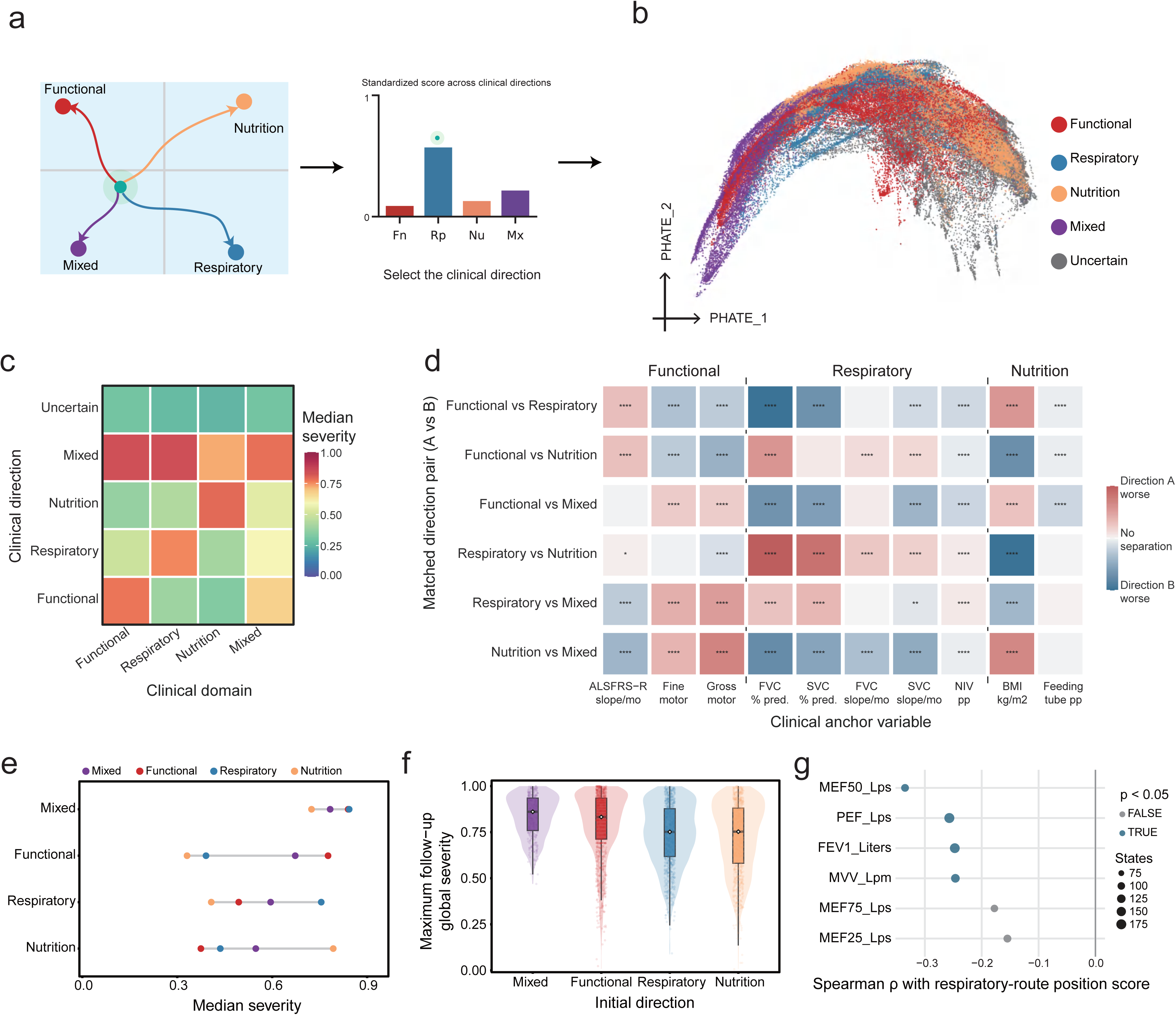
DynaALS resolves clinically distinct directions of deterioration. a. Four directions from a low-burden basin towards functional, respiratory, nutritional or mixed high-burden regions, assigned using graph-smoothed endpoint affinities. States with a maximum assignment score below 0.40 or normalized assignment entropy in the highest decile remained uncertain. b. PHATE display of the four confidently assigned directions and uncertain states; direction-specific state and participant counts are provided in Supplementary Table S5a. c. Median domain severity by assignment on a 0-1 scale; mixed states show multidomain burden. Values and interquartile ranges are provided in Supplementary Table S5a. d. Pairwise clinical contrasts after nearest-neighbour matching within cohort, burden quintile, ALSFRS-R quartile and reference-day bin. Same-patient pairs were excluded, although patients could contribute multiple states. Cell colour shows the direction-standardised paired difference; asterisks show BH-adjusted q values. Confidence intervals used 500 matched-pair bootstrap replicates; * q < 0.05, ** q < 0.01, *** q < 0.001 and **** q < 0.0001; ns, not significant. The complete contrast matrix is provided in Supplementary Table S5b. e. Median global and domain severity across 92,801 confidently assigned patient-time states; uncertain states were excluded. The four direction groups contained 11,963-30,722 states from 3,224-7,601 patients. Full summaries are provided in Supplementary Table S5d. f. Maximum global severity recorded from the initial state through available follow-up, stratified by the clinical direction of each patient’s earliest eligible state (n = 10,763). Multiple states on the same daywere deduplicated by prioritising ALSFRS-R landmarks, followed by fixed landmarks and final-observed states; uncertain initial directions were excluded. Mixed, functional, respiratory and nutritional initial directions contained 322, 3,007, 3,168 and 4,266 patients, respectively. Violin and box plots use all retained patients, whereas jittered points are a seeded display sample of up to 900 patients per direction. White diamonds denote medians. Direction-specific summaries are provided in Supplementary Table S5e. g. State-level Spearman associations in the supplementary AskHelpU pulmonary-function analysis, comprising states without contemporaneous standard FVC or SVC but with granular pulmonary-function measurements within 90 days. Exact correlations and nominal P values are provided in Supplementary Table S5g. Blue denotes nominal P < 0.05.

Direction added clinical composition to the global severity axis (Fig. 3c). Functional, respiratory and nutritional states were most severe in their corresponding domains, whereas mixed states carried high burden across all domains. These profiles persisted across the disease-burden range (Extended Data Fig. 7a). Functional states showed greater motor and total functional loss, respiratory states showed lower vital capacity, and nutritional states showed lower body-mass index. Mixed states accumulated all three forms of deterioration (Supplementary Table S5a,c). The atlas thus captured coordinated but nonuniform progression: several domains worsened together, while the dominant clinical direction differed between patient states. It also placed respiratory decline, linked to ventilatory support and mortality, and nutritional decline, linked to survival, within the same continuous geometry^24,25^.

Direction remained informative among states matched by cohort, global burden, ALSFRS-R and reference day (Fig. 3d). Fifty-one of 60 pairwise contrasts remained significant after Benjamini-Hochberg correction. At comparable overall burden, respiratory states had FVC values 18.9 percentage points below nutritional states. Nutritional states had a mean body-mass index 3.23 kg m^−2^ below respiratory states, and functional states declined 1.44 ALSFRS-R points per month faster than respiratory states. DynaALS therefore distinguished clinically different forms of deterioration among states with similar global severity and ALSFRS-R (Supplementary Table S5b).

Joint mapping of direction and severity revealed a multidomain convergence state alongside the three domain-dominant directions (Fig. 3e). Mixed states were severe across every domain. Respiratory severity was their strongest domain component, closely followed by functional severity, placing respiratory dysfunction prominently within the multidomain high-burden state (Supplementary Table S5d).

The earliest mapped direction also stratified the maximum global burden recorded from that state onward (Fig. 3f). Mixed initial states reached the highest median maximum severity (0.861). Among single-domain starts, functional states reached 0.832, compared with 0.752 for respiratory and 0.753 for nutritional states (Supplementary Table S5e). A functional-dominant initial state therefore marked a course that reached greater overall burden than respiratory- or nutritional-dominant starts. This linked the earliest clinical composition to the burden recorded across the subsequent patient timeline and extended earlier work on progression patterns and ordered impairment sequences^12,26^.

Directions remained coherent over months, while patient trajectories increasingly redistributed across them. Equal-cohort persistence decreased from 0.794 at 90 days to 0.666 at 180 days and 0.529 at 365 days (Extended Data Fig. 7b and Supplementary Table S5f). By one year, the equal-cohort mean placed about 47% of direction-assigned transitions in another direction. Increasing transitions into mixed and other directions showed how the dominant domain reorganised as ALS progressed. DynaALS therefore represents clinical direction as a changing property of the patient and follows how deterioration spreads across domains^11,12,26^.

AskHelpU further expanded this clinical resolution. Greater respiratory-direction position was associated with lower MEF50, PEF, FEV1 and MVV (rho = −0.335 to - 0.246; Fig. 3g and Supplementary Table S5g). Patient-contributed records thus reinforced the respiratory direction with pulmonary measurements beyond routinely harmonised vital capacity. Together, severity defined how far ALS had progressed, direction defined its current clinical composition, and serial remapping revealed how that composition changed over time.

### DynaALS predicts future-state geometry and decodes clinical outcomes

Earlier ALS models forecast aggregate ALSFRS-R decline from clinical-trial records^27^, simulated multiple functional impairments from an initial visit^15,16^, or predicted future assisted-ventilation risk from patient snapshots^28^. DynaALS extends these approaches by uniting clinical prediction with recovery of future disease-state geometry.

We first asked whether population-referenced transitions in DynaALS could recover subsequent disease states (Fig. 4a-d). Analogue-derived futures captured three complementary properties of progression: global severity, position within the atlas and clinical direction. Analogue-derived and observed future burden correlated at r = 0.888, 0.810 and 0.680 at approximately 90, 180 and 365 days, respectively (Fig. 4a). Observed futures were enriched 16.9- to 192.8-fold among the Top-10 to Top-30 analogue-supported candidate positions across these horizons (Fig. 4b). The observed future direction ranked first in 57.9%, 49.5% and 42.8% of states and within the top two in 78.4%, 72.0% and 65.0%, respectively (Fig. 4c). Leave-one-cohort-out analyses retained future-burden retrieval across all three resources and horizons (r = 0.647-0.886; Extended Data Fig. 8), supporting transfer between cohorts. Projection of three illustrative 180-day trajectories in PHATE showed the relationship between current, analogue-derived and observed future states (Fig. 4d).

**Fig. 4.**
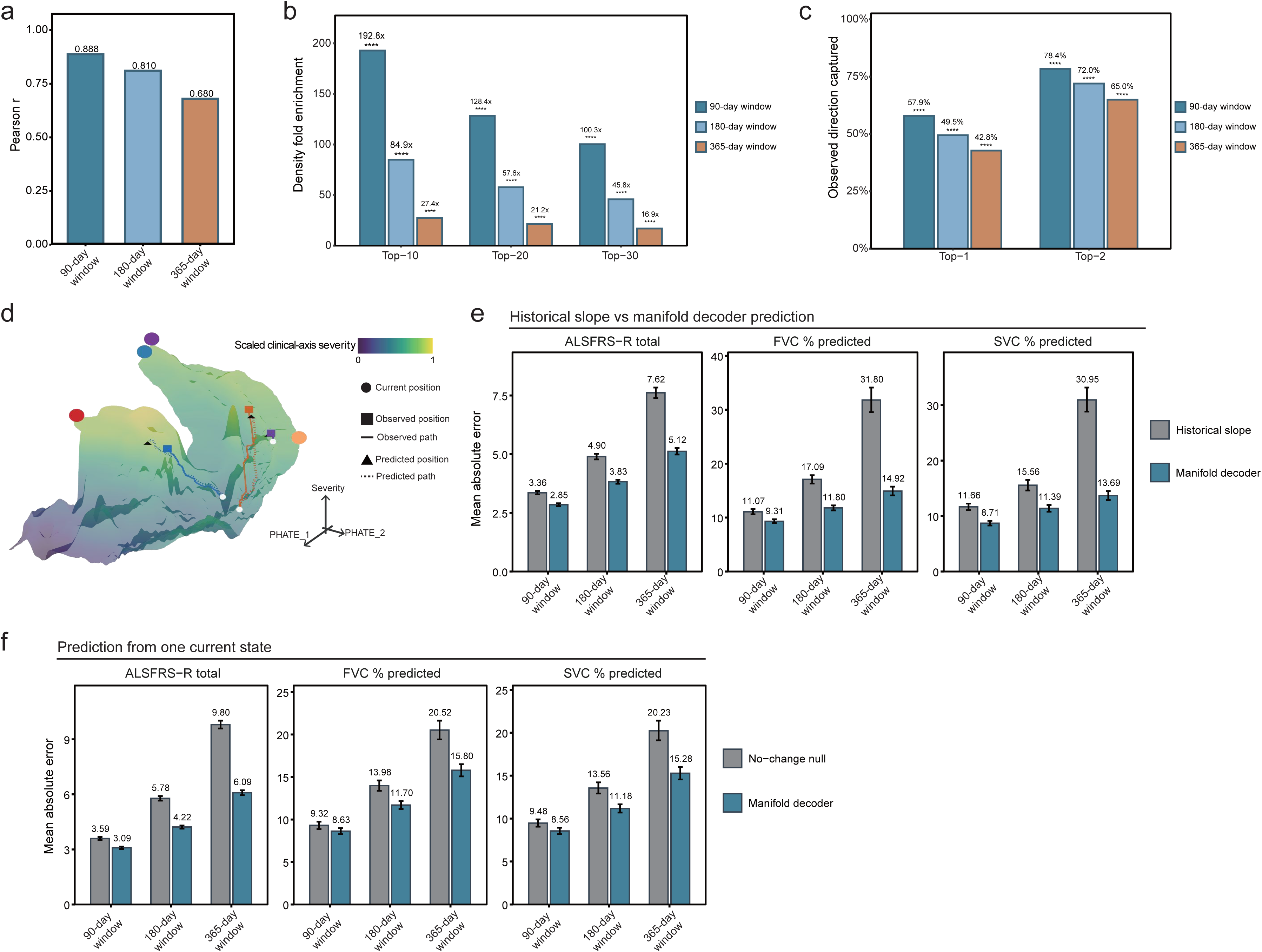
DynaALS forecasts future disease states and clinical outcomes. a. Observed versus analogue-derived future burden at 90, 180 and 365 days window. b. Fold enrichment of observed futures within Top-10, Top-20 or Top-30 candidate positions. Candidates included other patients’ futures plus the query future solely for ranking; other query-patient states were excluded. P values used one-sided hypergeometric tests; **** P < 0.0001. c. Top-1 and Top-2 capture of observed future direction against uniform 20% and 40% baselines for five possible labels (four clinical directions plus uncertain); P values used one-sided binomial tests; **** P < 0.0001. d. Three illustrative 180-day observed and analogue-derived paths shown with the low-burden basin and four high-burden regions in PHATE. e. ALSFRS-R, FVC and SVC errors at 90, 180 and 365 days for the DynaALS-conditioned future-state decoder versus bounded ordinary-least-squares slope extrapolation. Slopes required two same-target observations at least 30 days apart on identical complete cases. Patient-level out-of-fold ridge decoders used predicted future DynaALS coordinates, non-target-family future severity and clinical-direction summaries, and currently visible clinical covariates. Future-state construction excluded held-out patients from both transition edges and clinical-direction reference states (Supplementary Table S8a). f. Single-state, zero-history forecasting of ALSFRS-R, FVC and SVC from one current patient state. The future-state decoder was compared with a current-value no-change null. Bars show pooled MAE and 95% patient-cluster bootstrap confidence intervals from 5,000 replicates.

We then tested whether predicted future states could be decoded into quantitative clinical measurements. We then decoded predicted DynaALS states at 90, 180 and 365 days into future ALSFRS-R, FVC and SVC values, evaluating each participant with models fitted without that participant’s data. (Fig. 4e). Among participants with sufficient previous measurements to calculate a conventional progression slope, this approach produced smaller errors than slope-based extrapolation for all three outcomes at all three time points. Error reductions ranged from 15.2% to 55.8%, with the largest gains at 365 days. At 365 days, decoder versus slope MAE was 5.12 versus 7.62 ALSFRS-R points. FVC and SVC MAEs were 14.92 versus 31.80 and 13.69 versus 30.95 percentage points, respectively (Fig. 4e and Supplementary Table S8a).

We next tested whether forecasting remained possible from a single current state. This analysis used no earlier visits, so no patient-specific historical slope could be calculated (Fig. 4f). Using anchor-day information alone, the future-state decoder reduced pooled MAE relative to a current-value no-change null for all three outcomes at all three time points, with reductions of 7.4-37.9%. At 365 days, decoder versus no-change MAE was 6.09 versus 9.80 ALSFRS-R points. FVC and SVC MAEs were 15.80 versus 20.52 and 15.28 versus 20.23 percentage points, respectively (Supplementary Table S8b). Without any prior visits, single-state forecasts also yielded MAE point estimates 7.9-50.6% below the historical-slope errors in Fig. 4e for corresponding target-horizon cells. A single DynaALS state could therefore seed both future-state retrieval and multi-outcome clinical decoding without an individual longitudinal history.

MEDSTREM integrated these predictive functions with the record-construction workflow (Extended Data Fig. 13b). Once a longitudinal record had been reviewed, the application mapped its current state into DynaALS and displayed severity and clinical direction. It then forecast future states at 90, 180 and 365 days, decoded the corresponding clinical profiles and generated an LLM-assisted interpretation with follow-up questions. New observations could be incorporated to update the mapped state and forecasts within the same locally controlled workflow.

### DynaALS reveals clinically ordered molecular axes in motor neurons

We asked whether the multidimensional clinical geometry of DynaALS was also reflected in motor-neuron molecular variation. Answer ALS RNA-seq and ATAC-seq profiles were linked to each participant’s AIDI and four direction-peak coordinates (Fig. 5a,b). AIDI represented overall deterioration, while the four direction coordinates measured progression towards functional, respiratory, nutritional or mixed high-burden regions. Earlier Answer ALS studies showed that motor-neuron molecular profiles contain information about ALSFRS-R progression rate^10,18^. DynaALS extended this relationship from one scalar slope to the magnitude and clinical direction of deterioration.

**Fig. 5.**
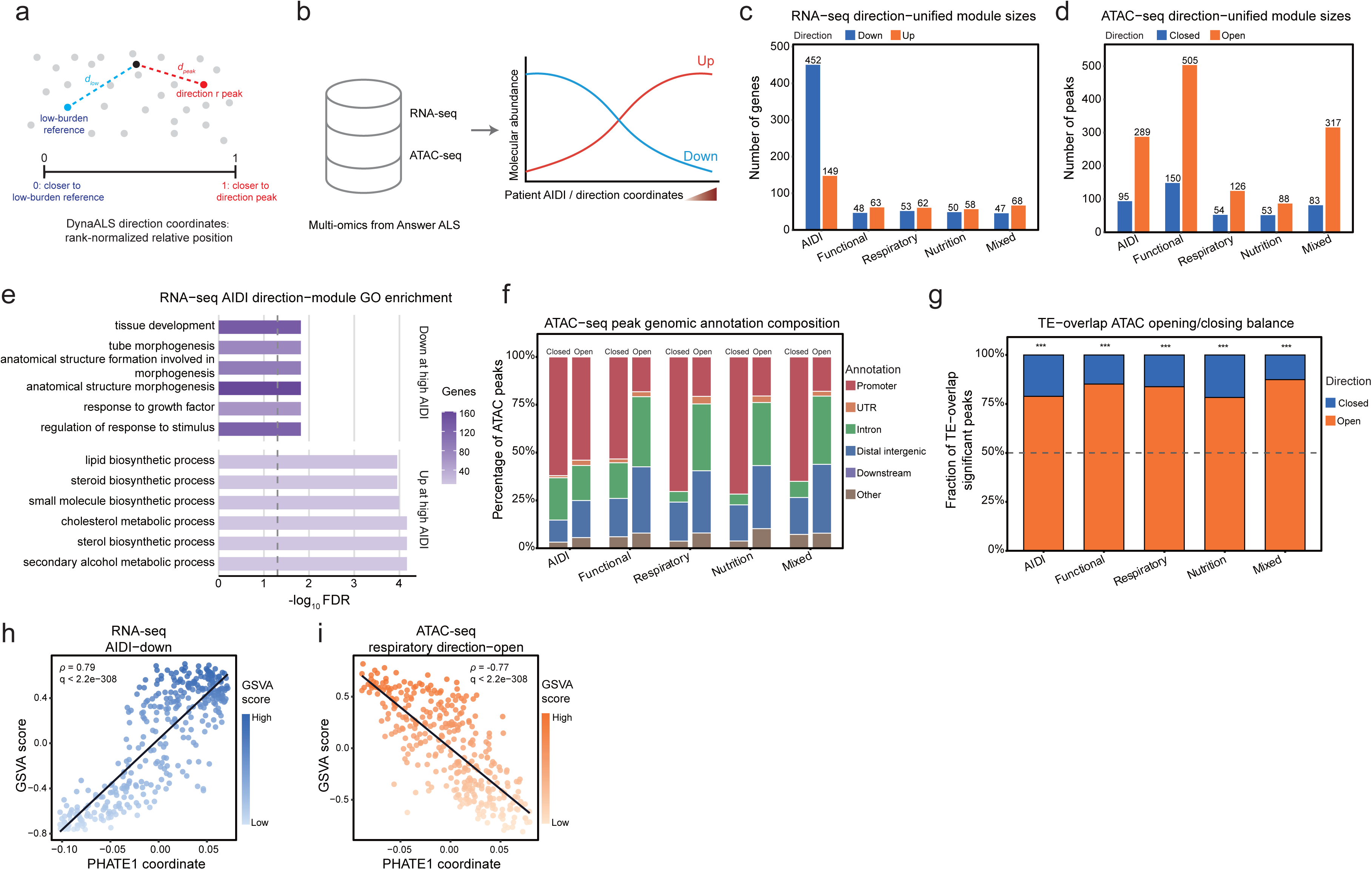
DynaALS organises motor-neuron RNA and chromatin programs. a. Patient-level direction-peak position used for molecular mapping. For each state and clinical direction, Euclidean distance to a low-burden centroid was divided by the sum of that distance and the distance to the nearest direction-specific high-burden peak centroid. The relative distance was rank-normalised across states and averaged across available states per participant; larger values indicate greater proximity to a direction-specific adverse peak. This coordinate is distinct from the graph-relative state position used in Fig. 3g. b. Answer ALS iPSC-derived motor neurons: RNA-seq, 475 patients; ATAC-seq, 455; paired, 428. Associations used AIDI percentile rank or four AIDI-adjusted direction-peak positions. c. Numbers of unique RNA genes at BH q < 0.05 from Monocle 2 negative-binomial spline likelihood-ratio tests. d. Numbers of unique ATAC-seq peaks at BH q < 0.05 from Monocle 2 negative-binomial spline likelihood-ratio tests; bars count features. AIDI-down and AIDI-up denote lower and higher expression at greater AIDI, respectively; AIDI-closed and AIDI-open denote lower and higher accessibility at greater AIDI. AIDI-adjusted direction signs in c,d used fitted high-minus-low contrasts with AIDI fixed at its median. e. Gene Ontology annotation of AIDI-down development/morphogenesis genes and AIDI-up cholesterol/sterol/lipid-biosynthesis genes. Bars show -log10(BH-adjusted P value); shading shows gene count. f. Genomic-annotation composition of selected ATAC peaks by accessibility direction; bars total 100%. Sets and open/closed assignments match in d. Closing regions contained a larger promoter-annotated fraction than opening regions, particularly across the four direction-associated sets. g. Transposable-element-overlap analysis of the AIDI set and four AIDI-adjusted direction peak sets (384, 655, 180, 141 and 400 selected peaks; 232, 445, 111, 83 and 278 transposable-element-overlapping peaks, respectively). Orange and blue denote higher and lower accessibility; two-sided binomial tests versus 50% were BH-adjusted across five quantities; ***q < 0.001 (all five comparisons). h. RNA PHATE1 association with the AIDI-down programme (n = 379, rho = 0.788, global BH q < 2.2 × 10-308). i. Partial Spearman association between AIDI-adjusted residuals of ATAC PHATE1 and the respiratory-opening programme GSVA score (n = 363, rho = −0.772, global BH q < 2.2 × 10-308). Analyses used one profile per patient. Feature counts and association metrics are provided in Supplementary Table S6a,b.

RNA expression and chromatin accessibility carried both levels of this organisation (Fig. 5c,d). AIDI defined broad molecular programs, and each clinical direction retained additional programs after adjustment for AIDI. Direction-associated RNA genes converged strongly across the four directions, whereas nearest-gene annotations of the corresponding chromatin sets were more direction-specific (Extended Data Fig. 9a,b). The clinical geometry therefore resolved a shared transcriptional response to worsening disease together with regulatory programs that distinguished how deterioration was expressed (Supplementary Tables S6a and S7d).

These molecular axes were biologically coherent. Greater AIDI was accompanied by lower expression of genes involved in development and morphogenesis and higher expression of cholesterol, sterol and lipid-biosynthesis genes (Fig. 5e). These changes accord with developmental, maturation and metabolic abnormalities reported in ALS iPSC-derived motor neurons^29–31^. After AIDI adjustment, all four directions retained a shared signature of chromatin organisation and developmental regulation. Functional-direction opening additionally highlighted neuronal differentiation and synaptic signalling (Extended Data Fig. 9c,d). This placed a common chromatin-organisation programme alongside a functional-direction neural-developmental branch, consistent with TDP-43-linked disruption of histone processing and chromatin structure^29,30,32^.

Chromatin direction was also evident in its regulatory context. Accessibility loss was concentrated near gene-proximal regulatory regions, whereas transposable-element-overlapping regions showed a consistent opening bias across the global and direction-associated peak sets (Fig. 5f,g). This pattern connects the DynaALS molecular axes to ALS progression signals, TDP-43-associated LINE decondensation and ALS-TE cortical programs related to TDP-43 dysfunction and disease duration^18,32–34^. DynaALS therefore revealed an ordered molecular disease space in which global deterioration and clinical direction were coupled to distinct transcriptional and regulatory programs.

Finally, we asked whether molecular variation learned independently from the omics data reproduced this clinical ordering. Unsupervised PHATE maps were fitted separately to RNA and ATAC profiles (Extended Data Fig. 9e,f). The AIDI-down RNA programme tracked RNA PHATE1 (rho = 0.788), and all four AIDI-adjusted direction-up RNA programs aligned with the same molecular axis (absolute rho = 0.800-0.825; Fig. 5h and Extended Data Fig. 9g,i). The global AIDI-open chromatin programme similarly tracked ATAC PHATE1 (rho = −0.775). After AIDI adjustment, all eight direction-associated opening and closing programs aligned with this axis (absolute rho = 0.670-0.861; Fig. 5i and Extended Data Fig. 9h,j). The molecular manifolds thus independently recovered both levels of clinical information captured by DynaALS: overall deterioration and direction-related structure beyond global burden (Supplementary Table S6b).

### DynaALS-linked programs emerge early during motor-neuron maturation

We next asked how early DynaALS-linked programs emerged during motor-neuron maturation. Using an established differentiation framework^35^, we generated iPSC-derived motor neurons (iMNs) from one control line and two ALS lines representing lower and higher AIDI. Each line was sampled on days 22, 27 and 33 for bulk RNA-seq and ATAC-seq, with three libraries generated per assay at each time point (Fig. 6a). Principal components separated line-timepoint combinations in both modalities (Fig. 6b). SMI32 staining documented iMN differentiation (Extended Data Fig. 10a), and transposable-element-associated RNA and chromatin features retained the line- and time-dependent structure (Extended Data Fig. 10b).

**Fig. 6.**
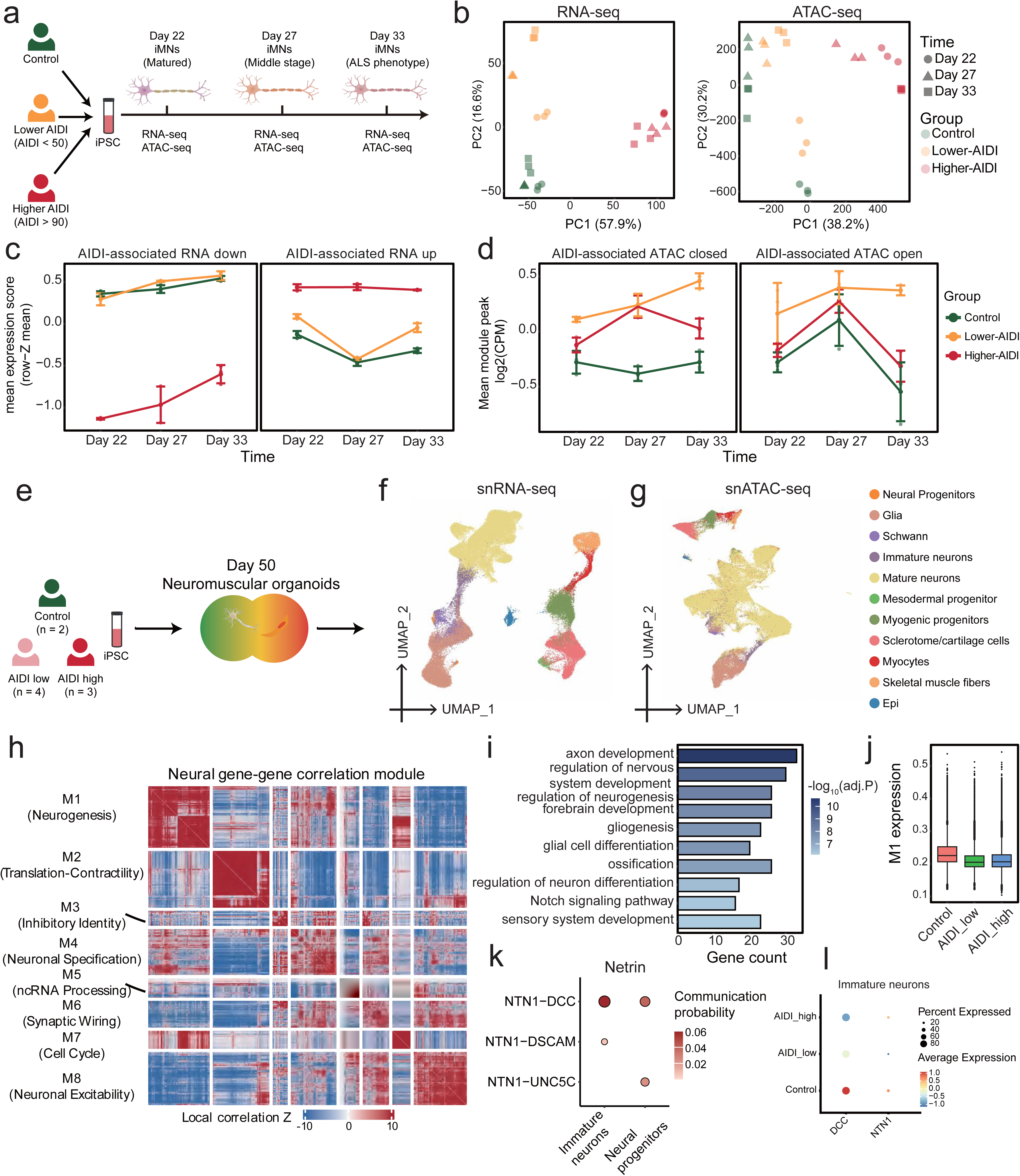
Patient-derived multi-omics links a neural-developmental module to a candidate Netrin-DCC axis. a. iPSC-derived motor neurons (iMNs) from one control line, one lower-AIDI ALS line (AIDI < 50) and one higher-AIDI ALS line (AIDI > 90). RNA-seq and ATAC-seq were collected on days 22, 27 and 33 with three replicate-labelled libraries per line-day and modality. b. Principal-component projections of 27 RNA-seq and 27 ATAC-seq profiles; colour shows line/group and shape shows day. c. AIDI-down/up RNA programs transferred across the iMN maturation time course without refitting. Points show line-day means and s.d. across three replicate-labelled libraries. d. AIDI-closed/open ATAC programs transferred across the iMN maturation time course without refitting. Points show line-day means and s.d. across three replicate-labelled libraries. e. Day-50 NMO design: two control, four lower-AIDI and three higher-AIDI samples. snRNA-seq labels comprised one pooled control, three lower-AIDI and four higher-AIDI ALS labels; snATAC-seq comprised one control and three labels per ALS group. f. Cell-type UMAPs of snRNA-seq. g. Cell-type UMAPs of snATAC-seq. h. NMO Hotspot gene-correlation Z-score matrix of eight co-expression modules (M1-M8). the complete programme-module map is shown in Extended Data Fig. 11c. i. Gene Ontology annotation of M1, a neural-development module containing NTN1 and DCC; bars show gene count and shading shows -log10(BH-adjusted P value). j. Cell-level M1 UCell scores across pooled control, lower-AIDI and higher-AIDI labels. k. CellChat-inferred NTN1-DCC, NTN1-DSCAM and NTN1-UNC5C interactions from Schwann cells to immature neurons or neural progenitors in the pooled 85,890-cell snRNA-seq atlas. l. Cell-level DCC and NTN1 DotPlot in immature neurons; size shows expressing-cell percentage and colour shows scaled mean expression.

Projection of the predefined Answer ALS programs into this time course revealed a coherent molecular ordering without refitting. The AIDI-down RNA programme was already lower in the higher-AIDI line on day 22 and remained separated from control on days 27 and 33 (Fig. 6c). AIDI-open chromatin followed the expected direction from the same early time point and across all three sampled days. AIDI-up RNA provided a parallel, lower-coverage signal, whereas AIDI-closed chromatin was less coherent (Fig. 6d). DynaALS-linked molecular ordering was therefore present at the earliest stage assayed, before the later maturation endpoint used by Answer ALS.

Persistent gene-level changes reinforced this cross-system continuity. Genes downregulated across all three time points overlapped more strongly between the lower- and higher-AIDI lines than persistently upregulated genes (Extended Data Fig. 10c,e). Shared losses converged on extracellular-matrix organisation, cell adhesion and wound healing. In the higher-AIDI line, they extended to DNA replication, chromosome segregation and cell-cycle programs (Extended Data Fig. 10f). Persistent increases highlighted embryonic patterning and axon guidance, linking the iMN time course to the developmental programs resolved below in NMOs (Extended Data Fig. 10d). Together, programme projection and persistent expression identified AIDI-down as the clearest transcriptional bridge from the Answer ALS clinical ordering to early iMN maturation. This convergence accords with early molecular abnormalities and patient-dependent emergence of cellular phenotypes reported in ALS iPSC-derived motor neurons^29,30,36^.

### Neuromuscular organoids single-cell multi-omics links a neural-developmental module to a candidate Netrin-DCC axis

To identify the cell types and interactions carrying these early programs, we profiled day-50 control, lower-AIDI and higher-AIDI NMOs by single-nucleus RNA sequencing (snRNA-seq) and single-nucleus ATAC sequencing (snATAC-seq) (Fig. 6e). Patient-derived NMOs reproduce motor-neuron, Schwann-cell and neuromuscular-junction pathology in C9orf72 ALS^37^, providing a multicellular setting for this analysis. Cell-type UMAPs resolved neural, glial and muscle compartments (Fig. 6f,g), and MyHC and alpha-bungarotoxin staining documented NMO generation (Extended Data Fig. 11a).

Within neural-lineage cells, Hotspot resolved eight local co-expression modules (M1-M8; Fig. 6h). The transferred AIDI-down programme aligned most strongly with M7 and next with M1, which were also the most positively coupled module pair (Extended Data Fig. 11b,c). M7 concentrated in neural progenitors and was enriched for nuclear division and chromosome segregation, recapitulating the higher-AIDI cell-cycle loss in iMNs. M1 was strongest in immature neurons and neural progenitors and was enriched for axon development, nervous-system development and synapse organisation (Fig. 6i and Extended Data Fig. 11d,g). M1 activity was lower in both ALS groups than in control (Fig. 6j). Transferred RNA and chromatin programs extended these DynaALS-linked signals across the neural-lineage landscape (Extended Data Fig. 11e,f,h). Module-level matrices and gene membership are provided in Supplementary Table S7a-c. The NMO atlas thus resolved the clinically anchored programme into a progenitor-associated proliferative branch and an immature-neuron neural-developmental branch.

The neural-developmental branch converged on Netrin-DCC signalling. NTN1 connected the Answer ALS AIDI-down programme, the persistent higher-AIDI iMN response, M1 and the CellChat ligand repertoire. DCC connected M1, the persistent response shared by both ALS iMN lines and the receptor repertoire (Extended Data Figs. 10c,e and 12a,b). CellChat placed this relationship in a multicellular context, inferring Schwann-cell-derived NTN1 signalling to immature neurons and neural progenitors through DCC, DSCAM or UNC5C (Fig. 6k and Extended Data Fig. 12c,d). NTN1 expression in Schwann cells and DCC expression across neural progenitors and neurons localised the predicted sender and receiver compartments (Extended Data Fig. 12e,f). Within immature neurons, NTN1 and DCC expression and DCC-locus accessibility were all lower in ALS NMOs than in control (Fig. 6l and Extended Data Fig. 12g). Together with reduced M1 activity, these clinical, temporal, single-cell, chromatin and communication layers converged on a DynaALS-linked neural-developmental Netrin-DCC axis. Prior studies independently implicated DCC-linked axon guidance in ALS: DCC was dysregulated in sporadic ALS iPSC-derived motor neurons with early axonal defects, and DCC silencing partially rescued neurite-complexity and growth-cone defects in UBQLN2-mutant motor neurons^38,39^. Our cross-scale analysis extends these observations by linking Netrin-DCC to DynaALS-defined clinical deterioration and multicellular neural-developmental programs, prioritising this axis for perturbational testing.

## DISCUSSION

This study establishes an end-to-end architecture for building dynamically updated patient-state representations from fragmented longitudinal evidence. MEDSTREM transforms patient-held medical-record images into locally controlled, reviewable histories. By combining these histories with established cohorts, DynaALS converts heterogeneous observations into a shared geometry of patient states. The resulting representation can be initialised from one current state, forecast through disease space, decoded into multidimensional clinical profiles and refreshed when new observations arrive. In ALS, this unifies capabilities that had remained split across data extraction, trajectory modelling, endpoint prediction and molecular association; more broadly, it offers a new model for turning distributed medical evidence into dynamic disease representations.

MEDSTREM changes both who can initiate ALS data construction and where computation occurs. Existing platforms and extraction systems have separately supported patient-mediated record aggregation, online clinical inquiry and conversion of medical documents into research data^40–45^. MEDSTREM brings these functions into one ALS workflow, linking patient-uploaded images, domain-tuned transcription, date-aligned extraction, reviewable personal records and cohort-scale analysis. Its fully local SuperOCR-Qwen3 workflow exceeded GPT-4o at each conversion stage under clean and light image corruption, and closely approached a DeepSeek V3-enabled workflow for structured extraction and longitudinal integration. MEDSTREM therefore couples competitive accuracy with privacy-preserving local control using workstation-scale models, allowing protected source documents to remain within authorised infrastructure^19–21^. By combining local processing with review, editing and export, MEDSTREM makes patient-held records available for personal record management and cohort-scale ALS research.

DynaALS changes the object of ALS modelling from a scalar outcome or patient-level subgroup to a repeated multidomain patient-time state. Earlier studies identified progression clusters, clinical subtypes, prognostic zones and ordered impairment sequences.^11–17,23,27,28^. DynaALS adds a time-resolved layer to these frameworks. AIDI measures the magnitude of deterioration, direction captures its clinical composition, and serial remapping follows its reorganisation. Burden-matched states retained distinct functional, respiratory and nutritional profiles, showing that similar global severity can arise through different clinical routes. Mixed states captured convergence across domains, with respiratory dysfunction prominent in the multidomain high-burden state^24,25^. I Functional-dominant initial states reached greater recorded global burden than respiratory- or nutritional-dominant starts, identifying an early route into broader deterioration. As persistence declined over one year, trajectories spread into mixed and alternative directions. This state-based geometry turns ALS heterogeneity into an observable within-patient dynamic: domains deteriorate together, but their relative dominance differs between patients and changes during progression^12,26^.

Prediction turns this geometry into a patient-state engine. Population transitions recovered future severity, atlas position and clinical direction, after which state-aware decoders translated predicted future states into multiple clinical measurements. Crucially, DynaALS could initialise these forecasts from a single current state without requiring an individual’s longitudinal history. This single-state, zero-history capability addresses the clinical cold-start setting in which patient-specific slopes cannot be estimated and extends endpoint-specific ALS prediction models^15–17,27,28^. The MEDSTREM application operationalises this cycle by carrying reviewed longitudinal records through DynaALS mapping, multi-horizon forecasting, clinical decoding and interactive interpretation (Extended Data Fig. 13a,b). As new records arrive, a patient’s state can be remapped and population-referenced forecasts refreshed without rebuilding the atlas. Together, MEDSTREM and DynaALS implement the core computational loop of an ALS clinical digital twin, spanning ingestion, state location, forecasting, decoding, interpretation and updating^46^. This evolving representation provides a foundation for a continuously updated virtual patient that retains an individual’s position within a population disease model while generating clinically interpretable futures.

DynaALS also revealed that motor-neuron molecular variation is clinically directed. Earlier Answer ALS studies linked RNA activity and chromatin accessibility to ALSFRS-R progression rate^10,18^. DynaALS resolves the clinical heterogeneity compressed by that single slope. Molecular-only PHATE maps aligned with global AIDI programs and retained strong direction-programme structure after AIDI adjustment. The resulting RNA and chromatin manifolds carried information about both the magnitude and clinical composition of deterioration. RNA convergence across directions pointed to a shared transcriptional response, whereas the more direction-specific chromatin sets suggested distinct regulatory implementations. Developmental and lipid programs, promoter-biased chromatin closing and transposable-element-associated opening connected this molecular geometry to recurrent features of ALS biology^29–34^. The AIDI-down programme transferred into a separately generated iMN maturation series and was evident at the earliest sampled stage^29,30,36,37^. NMO single-cell multi-omics then decomposed this clinically anchored programme into coupled proliferative and neural-developmental branches across interacting cell types^37^. The AIDI-down programme aligned most strongly with M7, a neural-progenitor module enriched for nuclear division and chromosome segregation. Together with the higher-AIDI-specific loss of programmes governing DNA replication, chromosome segregation and cell-cycle progression across the iMN time course, this convergence identifies a recurrent proliferative component of the DynaALS-linked molecular programme, coupled to the M1 neural-developmental branch. The neural-developmental branch converged on lower M1 activity, reduced NTN1 and DCC expression and reduced DCC-locus accessibility in ALS NMOs. Together with Schwann-to-neural communication maps, these changes placed Netrin-DCC within a cross-scale programme spanning patient progression, motor-neuron maturation and multicellular neural development^38,39^. DynaALS therefore turns clinical dynamics into a coordinate system for experimentally addressable biology.

By making fragmented longitudinal medicine computationally continuous, MEDSTREM and DynaALS connect privacy-preserving data construction to an evolving disease-state model that can be mapped, forecast, decoded and biologically interpreted. Prospective cohort-locked evaluation of the current retrospective atlas, together with larger donor panels and direct perturbation of Netrin-DCC, will extend its clinical calibration and biological validation. This transferable architecture offers a practical route towards clinical digital twins in ALS and other complex diseases.

## Methods

### Study design and data sources

We developed DynaALS, the ALS Disease Dynamics Atlas, by integrating longitudinal clinical observations from Answer ALS, AskHelpU and PRO-ACT.9,10 The atlas contained 158,300 patient-time states from 21,371 participants (Answer ALS, 1,016; AskHelpU, 8,307; PRO-ACT, 12,048). The participant-level ALS Integrated Deterioration Index (AIDI) fitting set contained 21,381 canonical records (Answer ALS, 1,017; AskHelpU, 8,316; PRO-ACT, 12,048). DynaALS state construction additionally required at least one dated, observed, model-ready non-event clinical measurement. One Answer ALS record and nine AskHelpU records carried endpoint or risk-set information for AIDI fitting but lacked such a measurement and therefore yielded no DynaALS state; PRO-ACT counts were identical between the two analysis populations.

AskHelpU longitudinal records combined patient-uploaded medical documents and self-reported ALSFRS-R assessments from a continuously expanding cohort of more than 8,000 participants. Analysis-specific eligibility yielded 8,307 AskHelpU participants for DynaALS and 8,316 for AIDI. Answer ALS and PRO-ACT contributed research-cohort and clinical-trial observations, respectively. All records used for modelling were deidentified and processed under resource-specific access controls. Participant-level clinical data, participant-level AIDI values and donor-level molecular data remained controlled; the public release contains only disclosure-reviewed aggregate results.

The study also included patient-derived molecular measurements. Answer ALS induced pluripotent stem cell-derived motor-neuron profiles comprised RNA sequencing (RNA-seq) from 475 patients and assay for transposase-accessible chromatin using sequencing (ATAC-seq) from 455 patients^10,18^. The neuromuscular organoid (NMO) experimental design comprised two control, four lower-AIDI and three higher-AIDI experimental samples. Following modality-specific processing, the single-nucleus RNA sequencing (snRNA-seq) object contained seven ALS analysis labels plus a control reference. The single-nucleus ATAC sequencing (snATAC-seq) object contained six ALS analysis labels plus a modality-specific control reference. Experimental samples, analysis labels, donors, patients and cells were treated as distinct inferential units.

### AskHelpU enrolment and longitudinal data collection

Participants registered and logged into the platform via the “AskHelpU” WeChat public-account or AskHelpU website (https://askhelpu.com). During enrolment they submitted proof of a confirmed ALS diagnosis alongside basic personal information, which was manually verified by platform staff to ensure data integrity and accuracy (ExtendedData Fig. 1a).

After that, participants could regularly upload diagnostic reports, medical records, laboratory results, genetic testing reports and biomarker measurements (for example, neurofilament light chain, NFL) in the form of images, text or voice messages, thereby enabling longitudinal tracking of disease progression (Extended Data Fig. 1b). They could also complete standardized clinical assessment instruments—including the ALSFRS-R, ROADS and modified Norris scales—directly within the platform (Extended Data Fig. 1c). The system issues monthly reminders to encourage consistent reporting, and patients participating in clinical or research projects can complete these assessments under the guidance of clinicians or study coordinators.

### MEDSTREM clinical-record processing

MEDSTREM was a locally deployed workflow for patient-mediated construction of longitudinal records from uploaded document images, extending prior medical-document extraction and privacy-preserving local language-model workflows^19,21,43–45^. The four stages were image-to-Markdown transcription, Markdown-to-structured-record extraction, within-patient longitudinal integration and database storage. SuperOCR combined conventional OCR guidance with a locally fine-tuned Qwen2.5-Omni-7B model trained on synthetically degraded clinical-document images; training and inference details are provided in Supplementary Methods S1. A batch implementation supported cohort-scale processing, while a single-user application enabled record upload, review, editing, longitudinal visualisation and export. Image transcription and downstream extraction were performed in an authorised local environment. The resulting records retained values, units, dates, provenance and source-quality flags.

Extraction fidelity was evaluated using a synthetic benchmark of 100 simulated patients with ALS and 20 documents per patient. Ground truth was specified independently at the image, document and integrated-record stages for each of four image-noise levels. The real-record generalisation check used 65-80 deidentified images per model. Image-to-text performance was assessed using character and word error rates, semantic similarity and structural fidelity. Structured extraction and longitudinal integration were evaluated using key-path accuracy after recursive flattening of predicted and reference records.

### Harmonised clinical observations and state landmarks

The analysis unit was a patient-time state, defined by one patient, one reference time and one cohort. Reference times comprised fixed follow-up landmarks, ALSFRS-R-supported landmarks and the final observed time point. Each state summarised observations near the reference time, during the preceding 180 days and across the full available history, thereby capturing both current status and longitudinal change.

Source-specific fields were mapped to canonical clinical variables before analysis. Supplementary Table S3d lists the clinical variables normalised for use in the current study, together with their resource-specific fields, units, derivations and applied normalisation.

We defined a 42-concept state vocabulary across the three resources, comprising shared cross-resource concepts and high-coverage resource-specific concepts spanning functional scales, respiratory measurements, nutritional status, motor and neurological examinations, vital signs and laboratory observations. Each concept generated current and historical values, availability, recency and recent-change summaries. Window-level observation density, domain coverage and data-quality descriptors yielded 417 state features. Death, ventilation, feeding support and related events were retained in a separate event ledger and excluded from disease-state inputs.

High-confidence reference states required recent observations, multidomain coverage and prespecified data-support and quality criteria detailed in Supplementary Methods S2. Representation learning used patient-grouped partitions, and the fitted representation was then applied consistently across downstream geometry and transition analyses.

### Calculation of AIDI

AIDI was a participant-level, percentile-scaled latent index of global deterioration. The fitted participant representation was

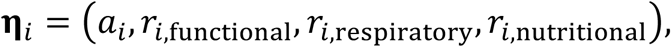

where *a_i_* captured deterioration shared across clinical domains and the three residual components absorbed domain-specific variation. This shared-plus-residual structure allowed AIDI to capture deterioration expressed across domains without forcing every endpoint onto a single trajectory. The reported index was the training-reference rank of the global latent:

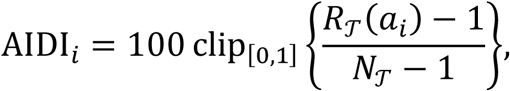

where *R*_*T*_(*a_i_*) is the average rank of *a_i_* relative to the pooled 70% training-reference population *T*, selected by cohort-stratified partitioning, and *N*_*T*_ is its size. Rank calibration removed dependence on the arbitrary numerical scale of the latent variable and yielded a bounded, interpretable index referenced to the training population^47^. Higher values therefore indicated greater global deterioration within the fitted population.

The latent measure was fitted jointly across the three resources using eight functional, respiratory and nutritional outcomes, including ALSFRS-R decline, FVC and SVC decline, weight loss and time to nutritional support. Grip strength, creatinine, creatine kinase, albumin and electromyography were reserved for post-fit association analyses.

Mortality status and event time were not used as endpoint-specific supervision. Cohort balance, endpoint availability and identifiability were incorporated during fitting, and the reported AIDI percentile was calibrated against the 70% training-reference population. Full model specifications are provided in Supplementary Methods S4.

### Clinical construct alignment

Clinical meaning was evaluated using four longitudinal outcomes: FVC percent-predicted decline, SVC percent-predicted decline, grip-strength decline and body-weight decline. For each outcome, AIDI was compared with ALSFRS-R decline rate and two descriptive full-follow-up summaries: a regional/support milestone-stage proxy and a functional-domain-loss-count proxy. These summaries were informed by established ALS staging concepts but were not treated as formally adjudicated King’s or MiToS stages^6–8^.

Each outcome analysis used the same complete-case participants for AIDI and all comparators. Associations used Spearman correlation after orienting measures towards deterioration. Cohort-stratified 5,000-replicate patient bootstrap estimated uncertainty and paired differences. Benjamini-Hochberg correction was applied within prespecified comparator families. FVC and SVC construct analyses used re-extracted, unit-verified percent-predicted measurements with valid dates and sufficient repetition. Missing grip values were not imputed.

### The dynamic disease manifold underlying DynaALS

DynaALS was learned using a partitioned encoder that separated longitudinal clinical status from patterns of clinical observation and data availability, including missingness, recency, coverage and data quality. This separation allowed observation patterns to inform data support without permitting resource-specific documentation practices to define the clinical geometry. AIDI and complementary global and domain-specific deterioration measures supervised the clinical representation, while reconstruction and separation objectives limited cohort and observation-process effects. Metric refinement on high-confidence reference states yielded the shared coordinates applied to all patient-time states. Longitudinal dynamics were represented as movement through this geometry, which was visualised using PHATE^48^.

For held-out states, functional, respiratory and nutritional severity was reconstructed from neighbouring high-confidence reference states drawn either from the same resource or from the other two resources. Clinical directions were identified by smoothing functional, respiratory, nutritional and mixed burden over the DynaALS neighbourhood graph, treating each direction as a locally coherent pattern rather than a label triggered by one measurement. Low-confidence states were retained as uncertain to preserve ambiguous boundary regions. A graph-relative coordinate placed progression towards each adverse direction on a common low-to-high burden scale; graph construction, assignment thresholds and sensitivity analyses are detailed in Supplementary Methods S3.

### Clinical directions and observed dynamics

Clinical directions summarised global, functional, respiratory and nutritional burden in DynaALS. We first profiled all confidently assigned patient-time states. For Fig. 3f, each participant was assigned the clinical direction of their earliest eligible state after same-day deduplication; ties on the same day prioritised ALSFRS-R landmarks, followed by fixed landmarks and final-observed states. Maximum global severity was calculated across that initial state and all later eligible observations, irrespective of subsequent direction. Separately, the observed transition matrices selected one prespecified source landmark per participant, prioritising the earliest eligible ALSFRS-R landmark and otherwise the earliest eligible state. Each source was paired with the nearest state in 90-, 180- and 365-day windows; analyses retained confident source and target directions and equally averaged resource-specific row proportions. Burden matching compared directions among states aligned for resource, global burden, ALSFRS-R and follow-up time; raw contrasts retained clinical units, whereas standardised contrasts supported comparison across variables measured on different scales^49^.

The supplementary pulmonary-function analysis in Fig. 3g included AskHelpU patient-time states without contemporaneous standard FVC or SVC but with at least one granular pulmonary-function measurement recorded within 90 days. Associations between graph-relative respiratory position and MEF50, MEF75, MEF25, PEF, FEV1 and MVV were assessed using state-level Spearman correlations. All six measurements are shown in Fig. 3g and reported in Supplementary Table S5g; nominal P values are reported because this exploratory analysis did not adjust for repeated observations or multiple pulmonary measurements.

To benchmark AIDI against an established ALSFRS-R trajectory-stratification algorithm, eligible longitudinal ALSFRS-R trajectories from each resource were assigned to the published mixture-of-Gaussian-process (MoGP) reference model^11^. Reference clusters were ordered by their stored mean annual decline and compared with cluster-level AIDI. Because both analyses contained ALSFRS-R information, this was interpreted as cross-method concordance.

### Near-horizon transition analogues

Near-horizon transition analogues were evaluated at approximately 90, 180 and 365 days window using visit-spacing tolerances, extending patient-similarity and time-window prediction strategies used in earlier medical and ALS models^27,28,46^. The analogue distance favoured transitions that began near the query, occurred over a compatible horizon and shared clinical burden and available prior movement, while down-weighting weakly supported records. For query state *q*, candidate transition *e* received the normalised weight

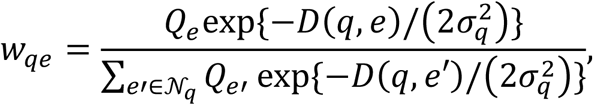

where *D*(*q*, *e*) combined source-state position, horizon agreement, available prior movement, current severity and data support, *Q_e_* was transition quality and *σ_q_* was the query-specific distance scale. The predicted future DynaALS position combined the weighted target positions with transport of the observed analogue displacements:

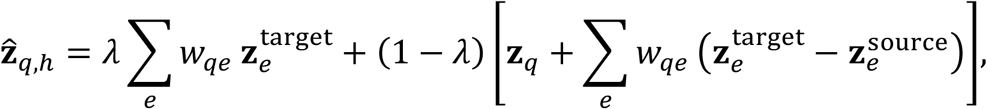

with *λ* = 0.5 in the selected model. The first term estimated a population-supported future destination, whereas the second transported the query from its current position using analogue displacements. Their blend balanced stability in absolute atlas position with preservation of query-anchored movement. The all-cohort regime used transitions from all cohorts, whereas the leave-one-cohort-out regime removed transitions from the entire target cohort. Query patients were excluded from the transition library in both regimes. For the illustrative paths in Fig. 4d, global and same-cohort forecasts generated by this operator were combined using the selected adaptive fusion.

Performance was summarised using correlations between observed and analogue-derived clinical burden, clinical-direction capture and future-state enrichment.

### Future clinical-variable decoding and single-state benchmarks

The core decoding targets were ALSFRS-R total, FVC percent predicted and SVC percent predicted at approximately 90, 180 and 365 days. For each held-out patient, transition analogues first projected the current state to horizon-specific future DynaALS coordinates, severity summaries and clinical-direction summaries. Patient-level cross-fitted ridge models then decoded the clinical target from this forecast state together with information available at the anchor. One head estimated the future value directly to stabilise the absolute clinical level, while a second estimated change from the current value to retain movement from the observed anchor. Their final prediction was

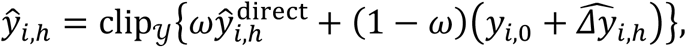

where *y* was the valid range of the clinical target. The ridge penalties and blend weight *ω* were selected separately for each target and horizon by nested patient-grouped cross-validation. No observations after the anchor were used. The snapshot and time-window information sets were aligned with prior ALS prediction designs.

For Fig. 4e, the latest eligible anchor was retained for each patient, target and horizon. Eligibility required at least two visible same-target observations spanning at least 30 days. A conventional comparator extrapolated the current value using an ordinary-least-squares monthly slope fitted to all visible same-target measurements. Both decoder and slope predictions were clipped to 0-48 for ALSFRS-R total and 0-200 for FVC or SVC percent predicted, and their performance was evaluated in identical complete cases. For Fig. 4f, only observations visible on the anchor day were admitted. Historical slopes, recent velocities and decline summaries were unavailable and encoded as missing; no earlier visit was used. The current-value no-change comparator was unfitted and carried the anchor-day raw target value forward.

Pearson confidence intervals used Fisher’s z approximation, and mean absolute error intervals used 5,000 patient-cluster bootstrap replicates (Supplementary Table S8a,b).

### DynaALS molecular programme mapping

The DynaALS molecular programme mapping shown in Fig. 5 used clinical quantities computed before molecular testing. In contrast to the graph-relative state positions used in Fig. 3, the four molecular direction-peak positions were derived from Euclidean distances in DynaALS to a low-burden centroid and the nearest direction-specific high-burden peak centroid. The resulting relative distance was rank-normalised across states and averaged within participant. RNA-seq and ATAC-seq associations with AIDI and these four patient-level direction-peak positions were tested using negative-binomial feature models and weighted standardised molecular summaries. Direction-peak associations were adjusted for AIDI and interpreted as molecular variation associated with clinical geometry beyond the global deterioration coordinate. Molecular measurements were not used to fit AIDI or DynaALS.

RNA-seq and ATAC-seq feature tests were corrected within prespecified multiplicity families. Selected ATAC-seq sets were summarised by genomic-annotation composition and overlap with RepeatMasker transposable-element intervals. These summaries described the selected regions and did not constitute genomic-background enrichment tests. Feature models, programme definitions and annotation procedures are provided in Supplementary Methods S7.

### iMN maturation time-course analysis

Induced pluripotent stem cells were maintained in mTeSR Plus on Matrigel-coated plates and passaged at least three times before differentiation. Sequential small-molecule induction generated motor-neuron progenitor spheroids, which were plated and matured with neurotrophic factors from day 16, building on established ALS iPSC-derived motor-neuron differentiation systems. The complete culture conditions are provided in Supplementary Methods S8.

The iMN maturation time course comprised motor neurons differentiated from one control iPSC line, one lower-AIDI ALS iPSC line and one higher-AIDI ALS iPSC line, sampled on differentiation days 22, 27 and 33. Three libraries per modality were available for each line and day. Metadata did not resolve whether these were biological or technical replicates; they were therefore treated as repeated within-line profiles rather than independent biological replicates.

RNA-seq and ATAC-seq principal-component analyses provided quality and trajectory views. Features were standardised across the 27 profiles and then averaged by line, day and programme to visualise within-line trajectories.

Parallel principal-component analyses restricted RNA to transposable-element-associated genes and ATAC-seq to transposable-element-overlapping peaks across all sampled days. Persistent line-associated genes were defined by BH-adjusted P < 0.05 and the same fold-change direction at each of the three days. Fisher’s exact tests quantified overlap between lower-AIDI-versus-control and higher-AIDI-versus-control persistent sets, and Gene Ontology analysis annotated the line-specific and shared partitions. Complete feature definitions and tested universes are provided in Supplementary Methods S8.

Temporal sensitivity was assessed using complementary programme-scoring, including gene set variation analysis, and ranked-feature analyses, with stability evaluated by leave-one-repeat-out analysis and persistence at later time points^50^. Because line identity was confounded with group, these analyses quantified programme-level temporal convergence. Complete thresholds are provided in Supplementary Methods S8.

### NMO single-cell multi-omics analysis

Neuromesodermal progenitors were induced for two days before aggregation in AggreWell400 plates, following a published ALS neuromuscular-organoid framework with study-specific modifications. NMOs were expanded under orbital culture and selected by bright-field morphology before day-50 molecular and imaging assays. The complete protocol is provided in Supplementary Methods S9.

The NMO design comprised two control, four lower-AIDI and three higher-AIDI samples. After modality-specific quality control, snRNA-seq and snATAC-seq resolved neural, glial and muscle populations. Hotspot identified eight co-expression modules (M1-M8), which were related to transferred DynaALS-linked programs using local-correlation scores and localised in neural-lineage cells with UCell^51^.

CellChat inferred ligand-receptor communication in the pooled snRNA-seq atlas, including Netrin-family signalling from Schwann cells to immature neurons and neural progenitors^52^. NTN1 and DCC expression and DCC-locus accessibility were then examined across annotated cell types. Cells and nuclei were treated as subsamples; experimental samples or analysis labels were the relevant biological units. Full processing, analysis-label composition and statistical specifications are provided in Supplementary Methods S9.

Cross-scale integration linked predefined Answer ALS programs to Hotspot modules, cell-level programme scores, pooled CellChat-inferred interactions and DCC-locus accessibility, connecting the neural-developmental M1 module to a candidate Netrin-DCC signalling axis for perturbational testing. Analysis units are detailed in Supplementary Methods S9.

### Statistical reporting and reproducibility

Continuous associations were reported with effect sizes, confidence intervals or exact permutation probabilities, as appropriate. Multiple testing was controlled using Benjamini-Hochberg correction within prespecified families. Patient-level, donor-level and cell-level units were distinguished explicitly, and cells or nuclei were never treated as independent biological replicates. Complete tested universes, exclusion rules, quality-control records, parameter manifests and software-session information for all primary analyses were retained in the controlled analysis package.

The public release contains aggregate source tables, plotting code, synthetic interface tests and disclosure-reviewed documentation. Fitted model parameters, patient-level clinical data, patient-level molecular summaries and donor-level NMO data remain in controlled environments.

## Data availability

The raw data generated in this study are available via the BioProject service of the China National Center for Bioinformation (https://www.cncb.ac.cn/) under accession number PRJCA050199. De-identified clinical data are available from AskHelpU (https://www.askhelpu.com/) under a data-use agreement. Details on how to request access to these clinical data are provided at https://www.askhelpu.com/?DataSharingStatement.

## Code availability

The code used for clinical and omics data analysis and visualization is provided in the Supplementary Data. De-identified example clinical report images are available in the Zenodo. A de-identified MongoDB sample dataset is provided as the_first_batch.gz and can be restored and inspected with MongoDB Database Tools using the command “mongorestore --gzip --archive=the_first_batch.gz”.

## Ethics Declaration

## Ethics statement

This study was approved by the Hangzhou First People’s Hospital review board (IIT-20240116-0012-02). All participants were fully informed about the study and provided written consent.

## Acknowledgements

We thank members of AskHelpU for providing supports in data collection. We thank members of the Tian and Liu laboratories for helpful discussions. This work was supported by the National Natural Science Foundation of China (grant nos. 32370784 and 22DAA01467), the National Key R&D Program of China (grant no. 2022YFA1105700 and 2022YFC3400400), the Westlake Education Foundation (WU2024WF002) and the Major Project of Guangzhou National Laboratory (grant nos. GZNL2024A01015 and YW-YFYJ0301).

## Author contributions

L.T., X.L., F.Y. and L.C. designed and supervised the study. L.C. established the AskHelpU platform. Z.L performed data pre-process with help from Y.C, Y.F. and G.L., C.G and YuT.F. performed iMNs and NMOs experiments with help from H.Z, S.W, S.J and J.Y. J.K. constructed SuperOCR and helped Z.L. performed data analysis. Z.L. performed the majority data analysis and prepared figures. L.T., Z.L., C.G. and YuT.F. wrote the manuscript with the contributions from all the authors.

## Extended Data Figures Legends

**Extended Data Fig. 1.**
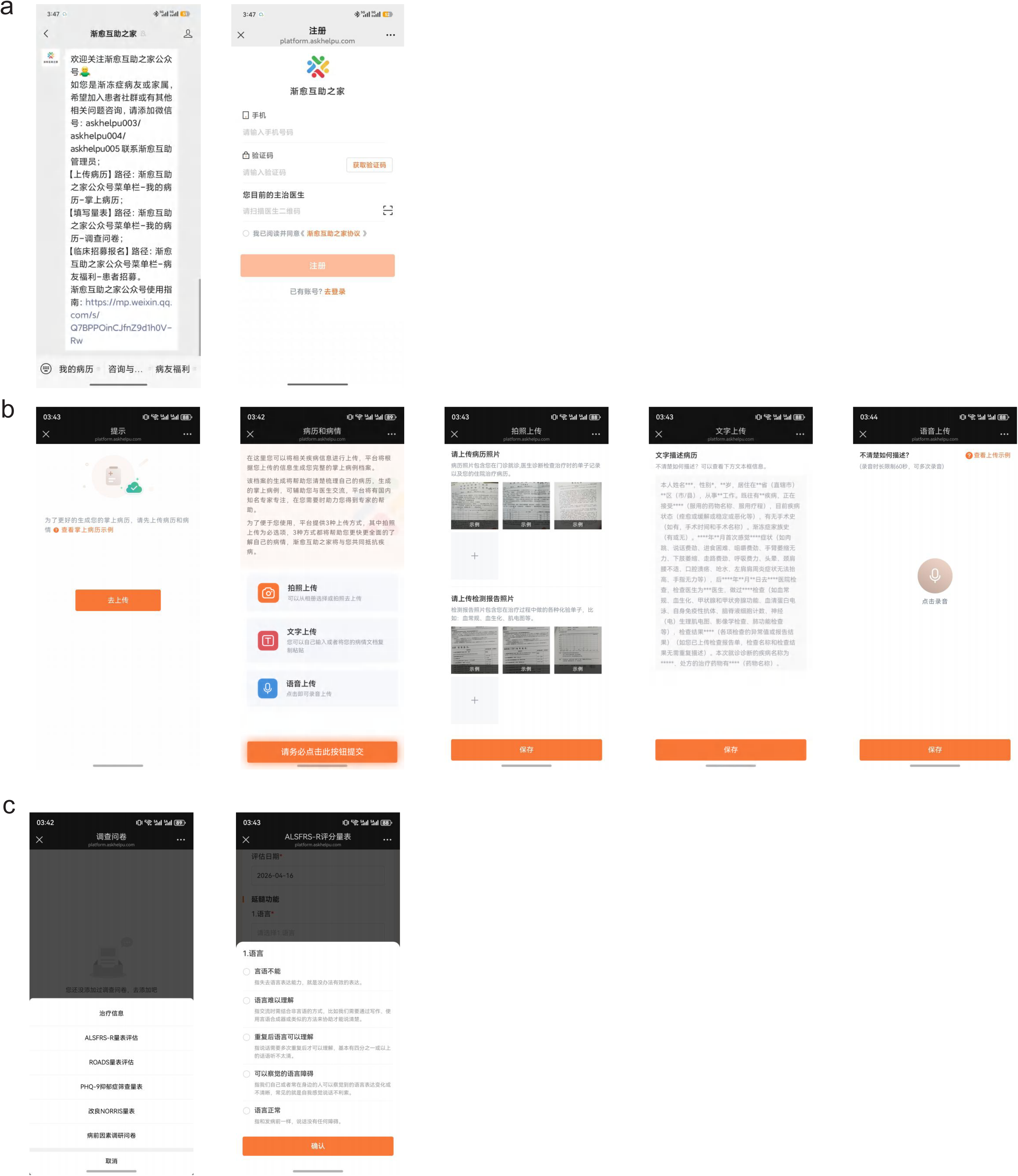
AskHelpU patient registration and longitudinal record collection. a. Registration through the AskHelpU WeChat service, including identity submission and manual review. b. Medical-record upload interface accepting existing files, camera images, typed text and voice input. c. Patient-reported interface for repeated ALSFRS-R, ROADS and related functional questionnaires.

**Extended Data Fig. 2.**
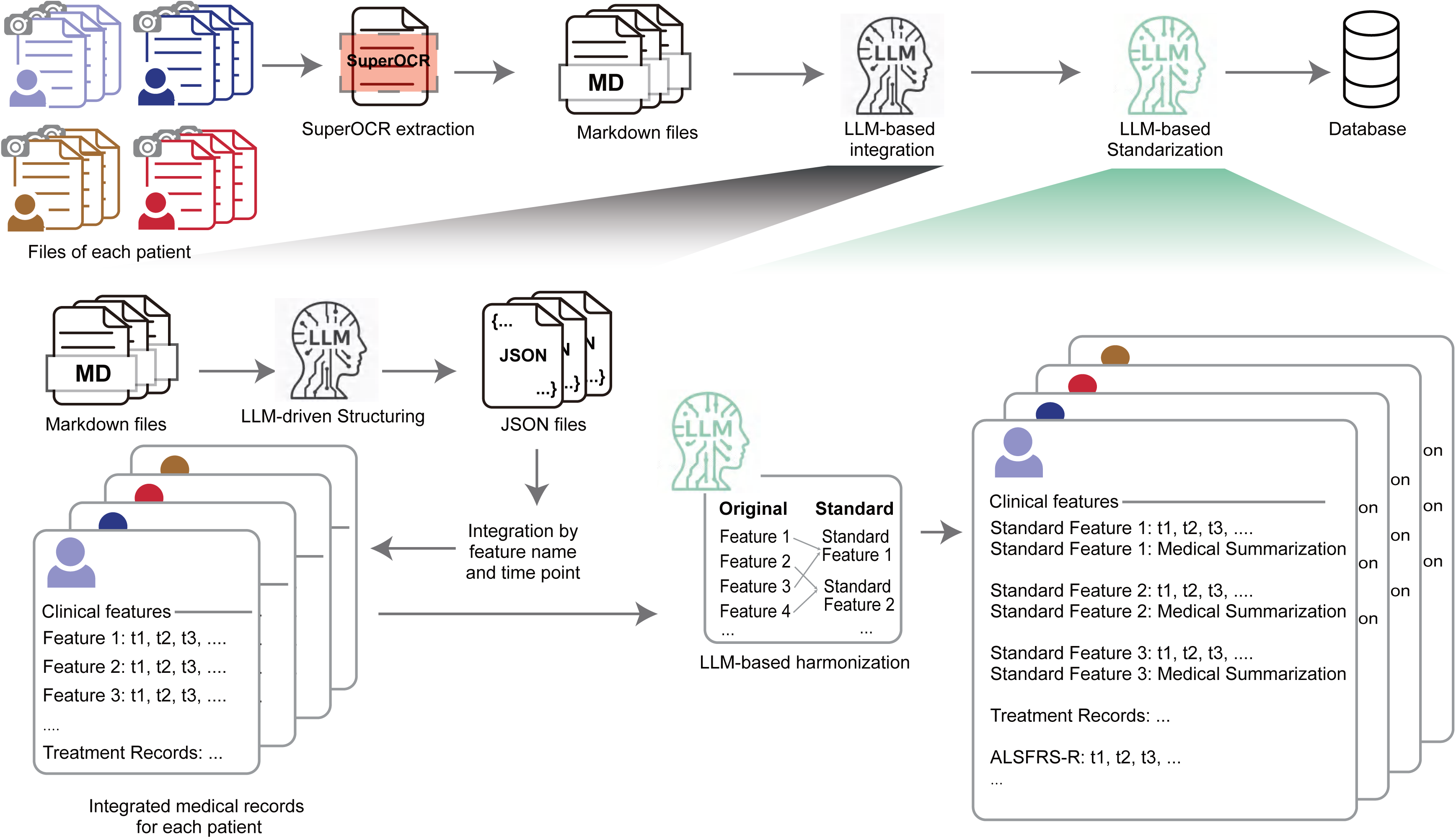
MEDSTREM processing and longitudinal-record harmonisation. Schematic of the MEDSTREM record-processing workflow. SuperOCR transcribed patient-level collections of uploaded medical-record images into Markdown. Large-language-model (LLM)-driven structuring converted each document into JSON. Dated document-level features were integrated into one longitudinal record per patient. An LLM-assisted terminology step mapped source features to standardized clinical concepts before database storage.

**Extended Data Fig. 3.**
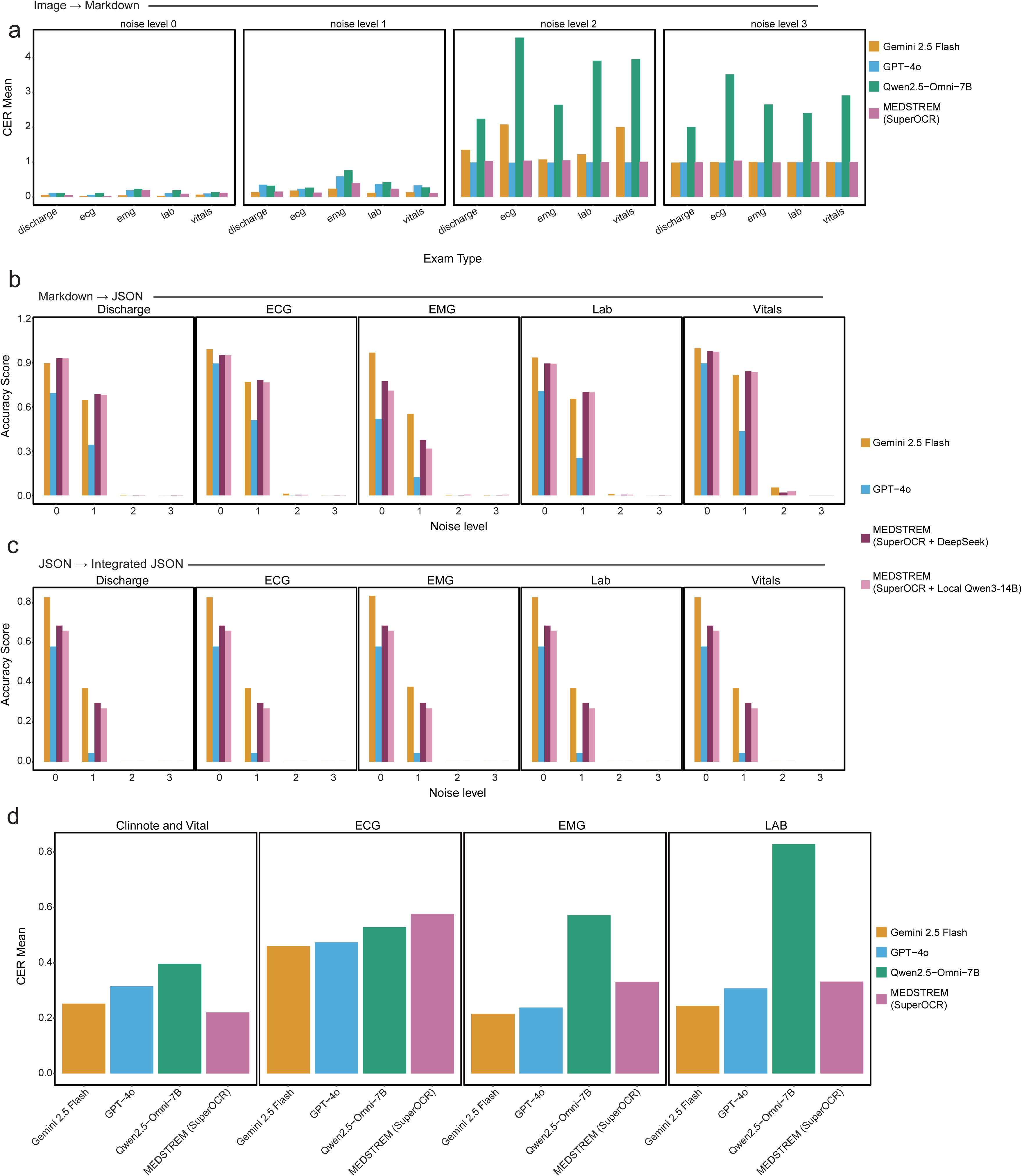
Stage- and document-type-resolved MEDSTREM benchmark. a. Mean character error rate (CER; lower is better) for synthetic discharge, electrocardiography (ECG), electromyography (EMG), laboratory and vital-sign documents across prespecified image-noise levels 0-3. Gemini 2.5 Flash, GPT-4o, Qwen2.5-Omni-7B and MEDSTREM SuperOCR were evaluated on the same synthetic document strata. b. Mean flattened key-path field accuracy (higher is better) for conversion of model-produced Markdown into document JSON by document type and noise level. Systems were Gemini 2.5 Flash, GPT-4o, MEDSTREM with DeepSeek V3 and MEDSTREM with local Qwen3-14B. c. Mean patient-level key-path accuracy after date-aware integration of model-produced document JSON, shown within the same document-type and noise strata. Integration was evaluated at patient level; repeated document-type facets are inherited display strata. d. Mean CER on manually deidentified real clinic-note or vital-sign, ECG, EMG and laboratory images for the four image-to-Markdown systems. Values are arithmetic means from the prespecified benchmark. The evaluations in b,c are chained and include propagated upstream errors, not independent tests beginning with ground-truth intermediate files. The underlying values are provided in Supplementary Table S2.

**Extended Data Fig. 4.**
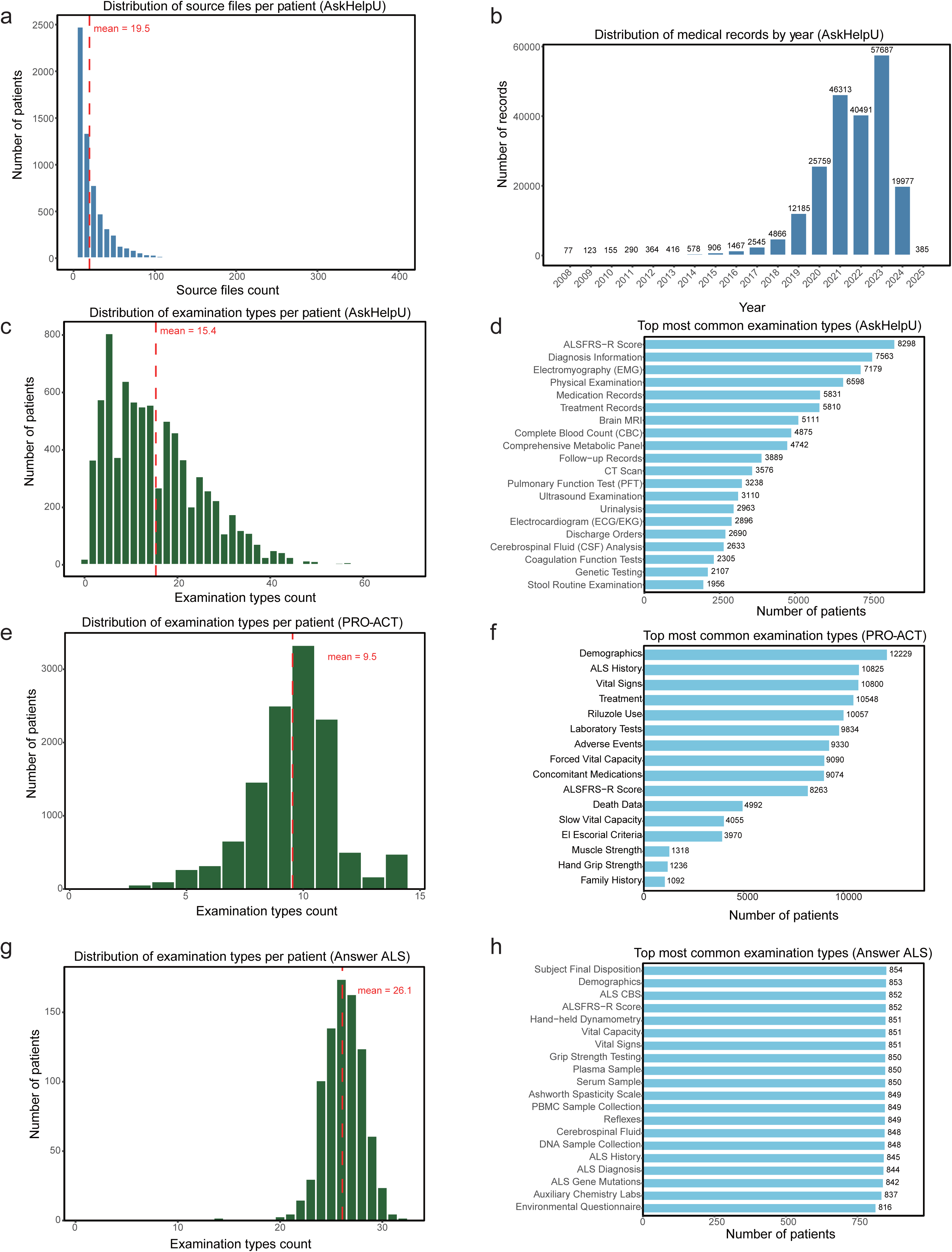
Source-data coverage before analysis-specific eligibility filtering. a. Distribution of source-file counts among AskHelpU participants retained in the file-coverage audit; the dashed line marks the mean of 19.5 files. b. Calendar-year distribution of AskHelpU medical records. c. Distribution of distinct examination-type counts among participants in AskHelpU. d. Ranked patient counts for most common examination tyles in AskHelpU. e. Distribution of distinct examination-type counts among participants in PRO-ACT. f. Ranked patient counts for most common examination tyles in PRO-ACT. g. Distribution of distinct examination-type counts among participants in Answer ALS. h. Ranked patient counts for most common examination tyles in Answer ALS. The underlying values are provided in Supplementary Table S3a-c.

**Extended Data Fig. 5.**
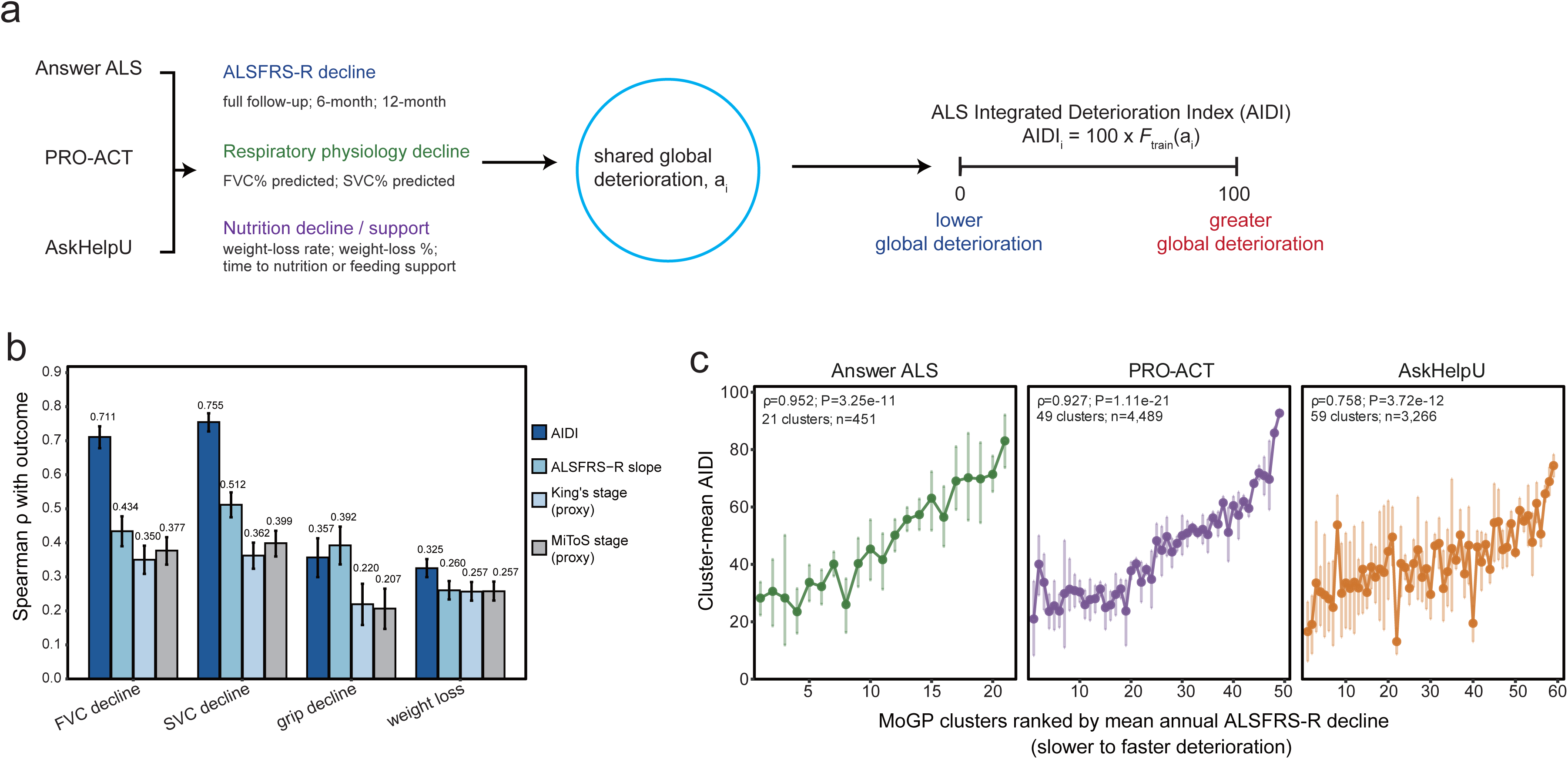
Clinical construct alignment of the ALS Integrated Deterioration Index (AIDI) a. AIDI definition. The index was estimated in 21,381 canonical participant records (Answer ALS, 1,017; PRO-ACT, 12,048; AskHelpU, 8,316). DynaALS contained 21,371 participants because state construction additionally required at least one dated, observed, model-ready non-event clinical measurement. One Answer ALS and nine AskHelpU records carried endpoint or risk-set information for AIDI fitting but lacked such a measurement and therefore generated no DynaALS state. A global deterioration latent trait was fitted jointly across the three resources using eight data-bearing functional, respiratory and nutritional outcomes; mortality status and time were not used for endpoint-specific supervision. The trait was converted to a 0-100 percentile using its empirical cumulative distribution in the 70% training-reference population. Higher AIDI indicates greater global deterioration over full follow-up; it is not a baseline prognosis or mortality-risk score. b. Same-patient complete-case Spearman correlations of AIDI, overall ALSFRS-R decline rate, a regional/support milestone-stage proxy and a functional-domain-loss-count proxy with FVC decline, SVC decline, grip-strength decline and weight loss. Outcomes were oriented so that larger values indicate greater deterioration, and each outcome used the same patient set for all four measures (n = 1,027-4,158). Error bars show 95% confidence intervals from 5,000 cohort-stratified patient-bootstrap replicates. c. Mean AIDI across published mixture-of-Gaussian-process (MoGP) reference clusters after cohort-specific assignment of eligible ALSFRS-R trajectories. Clusters were ordered by their stored mean annual ALSFRS-R decline. Points show cluster means; error bars show 95% confidence intervals from 5,000 within-cluster patient-bootstrap samples.

**Extended Data Fig. 6.**
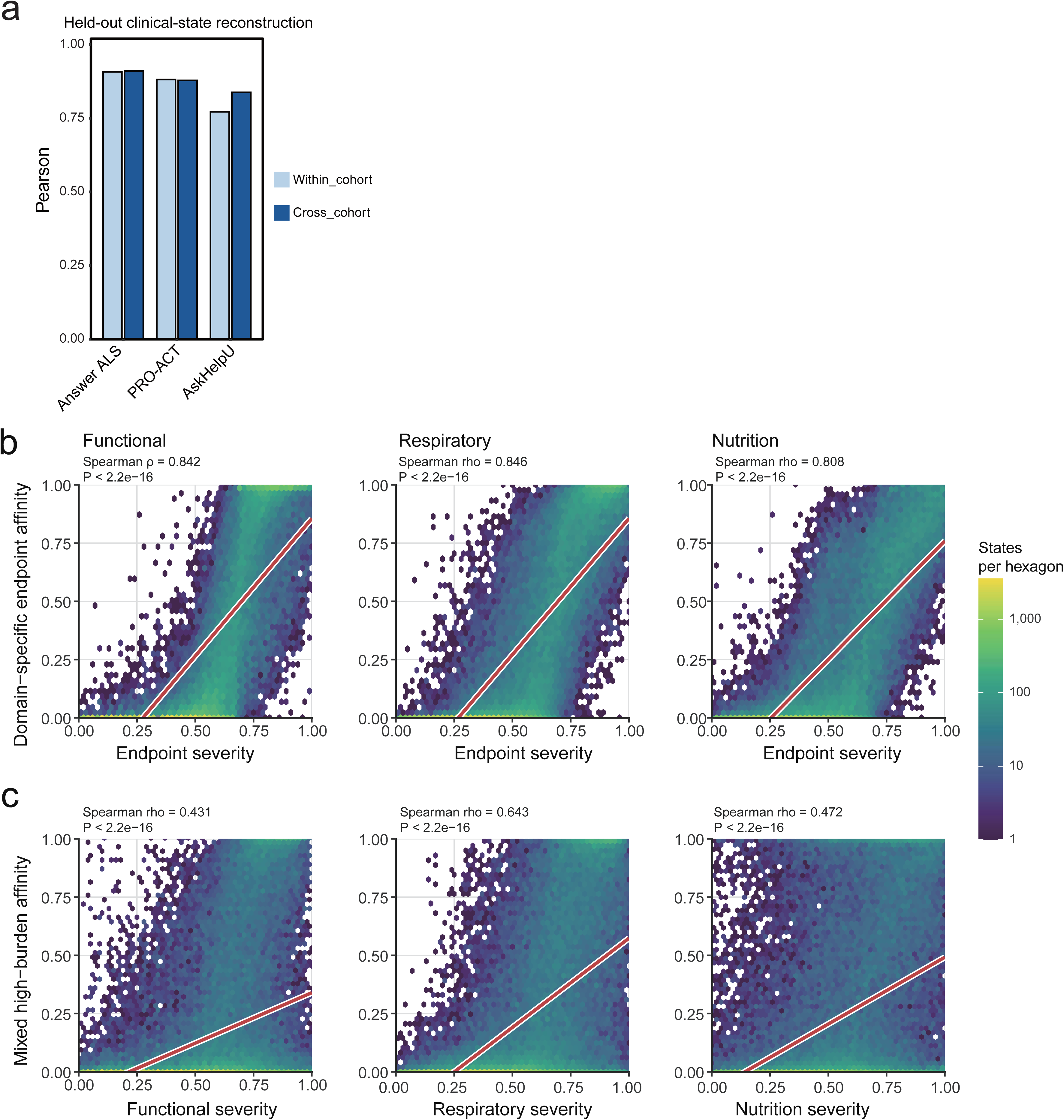
Reconstruction and clinical alignment of DynaALS. a. Reconstruction of functional, respiratory and nutritional severity in held-out context-test states by inverse-distance weighting of the 15 nearest high-confidence reference states in DynaALS. Within-resource reconstruction used reference states from the same resource, whereas cross-resource reconstruction used reference states from the other two resources. Bars show mean Pearson correlation between reconstructed and observed domain-severity targets for Answer ALS, PRO-ACT and AskHelpU. Within-versus cross-resource correlations are 0.908 versus 0.910 in Answer ALS, 0.882 versus 0.879 in PRO-ACT and 0.772 versus 0.838 in AskHelpU. b. Association between graph-smoothed DynaALS affinity and observed severity in the corresponding functional, respiratory or nutritional endpoint family across all patient-time states. Hexagons show state counts on a logarithmic colour scale. Red lines are display fits; labels report Spearman rho (0.842, 0.846 and 0.808, respectively; all P < 2.2 × 10^−16^). c. Association of mixed high-burden affinity with observed functional, respiratory and nutritional severity (Spearman ρ = 0.431, 0.643 and 0.472; all P < 2.2 × 10-^16^). Affinity was computed on the DynaALS graph, and endpoint scores are contemporaneous burden summaries.

**Extended Data Fig. 7.**
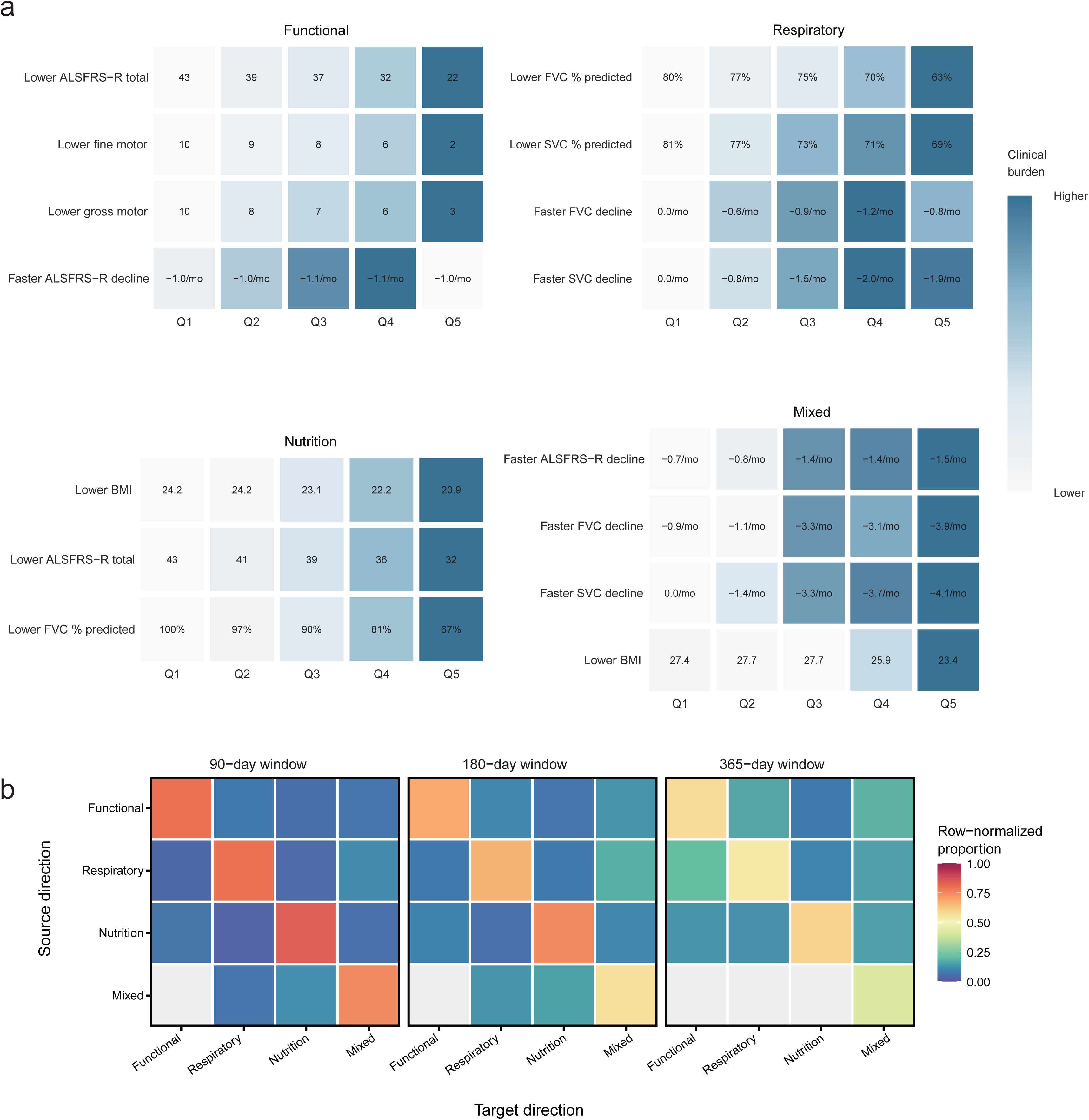
Direction-stratified burden and observed direction dynamics. a. Raw clinical measurements after stratification by functional, respiratory, nutritional or mixed clinical direction and by current global-burden quintiles Q1-Q5 computed over the full master state set. Cells show the observed median for each direction and quintile; colour is scaled from the least to the most adverse value separately within each clinical row and cannot be compared between variables. At greater burden, functional-direction quintiles show lower ALSFRS-R total, fine-motor and gross-motor scores. Respiratory-direction quintiles show lower FVC and SVC percentage predicted and faster decline. Nutritional-direction quintiles show lower BMI, whereas the mixed direction combines worsening functional, respiratory and nutritional measures. Quintiles are based on current global burden, not graph-relative direction position or AIDI. The underlying values are provided in Supplementary Table S5c. b. Observed transitions among confident functional, respiratory, nutritional and mixed directions at 90, 180 and 365 days window.

**Extended Data Fig. 8.**
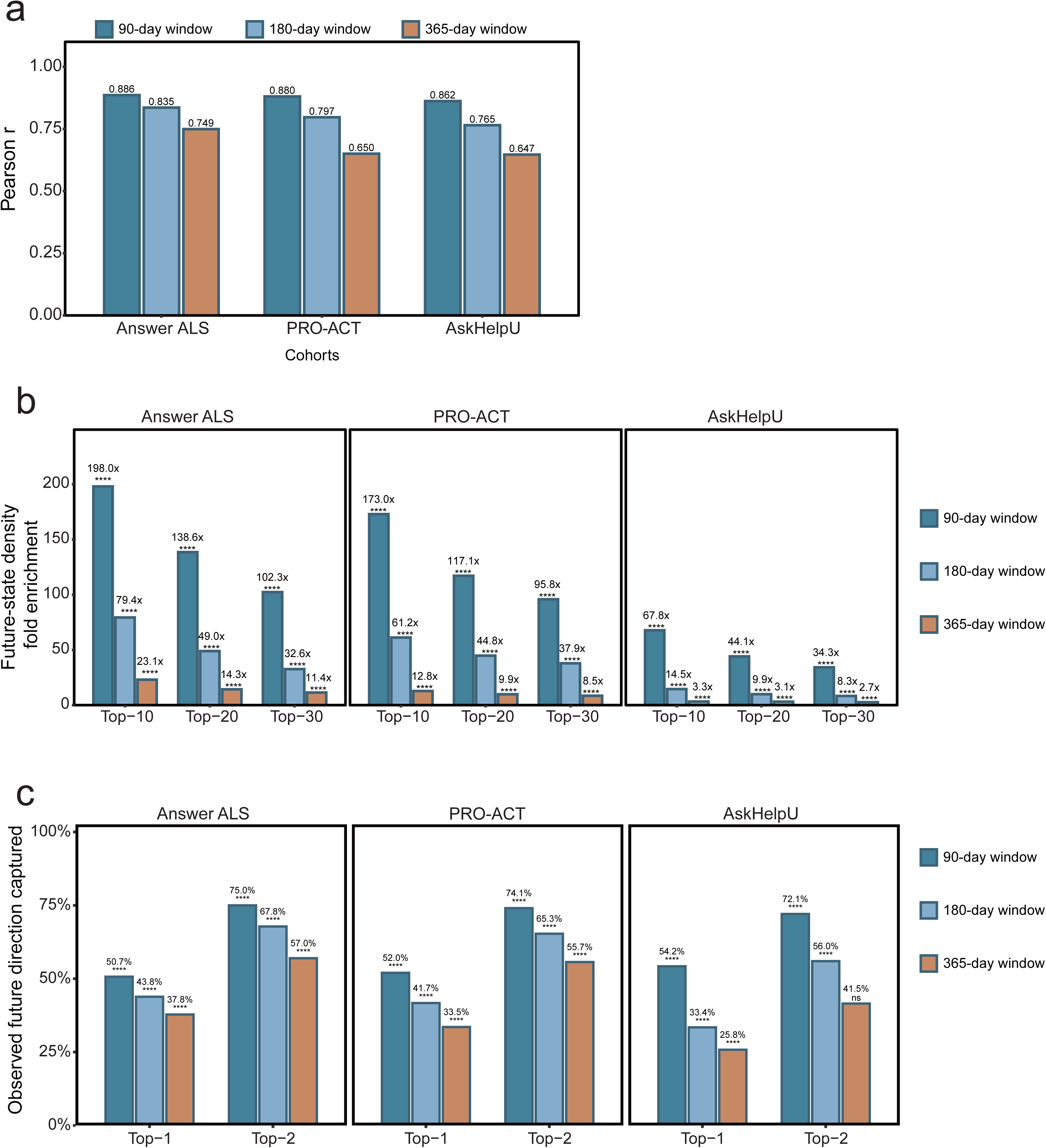
Leave-one-cohort-out evaluation of near-horizon transition analogues. a. Pearson correlation between observed and analogue-derived future clinical burden at approximately 90, 180 and 365 days when the target cohort was excluded from the transition library. b. Fold enrichment of observed-future-state density within the Top-10, Top-20 or Top-30 analogue-supported candidate positions relative to density outside those positions. P values are from one-sided hypergeometric tests; **** P < 0.0001. c. Top-1 and Top-2 capture of the observed future clinical direction. Dashed lines show uniform random baselines of 20% and 40% across five labels (four clinical directions plus uncertain); P values are from one-sided binomial tests; **** P < 0.0001; ns, not significant.

**Extended Data Fig. 9.**
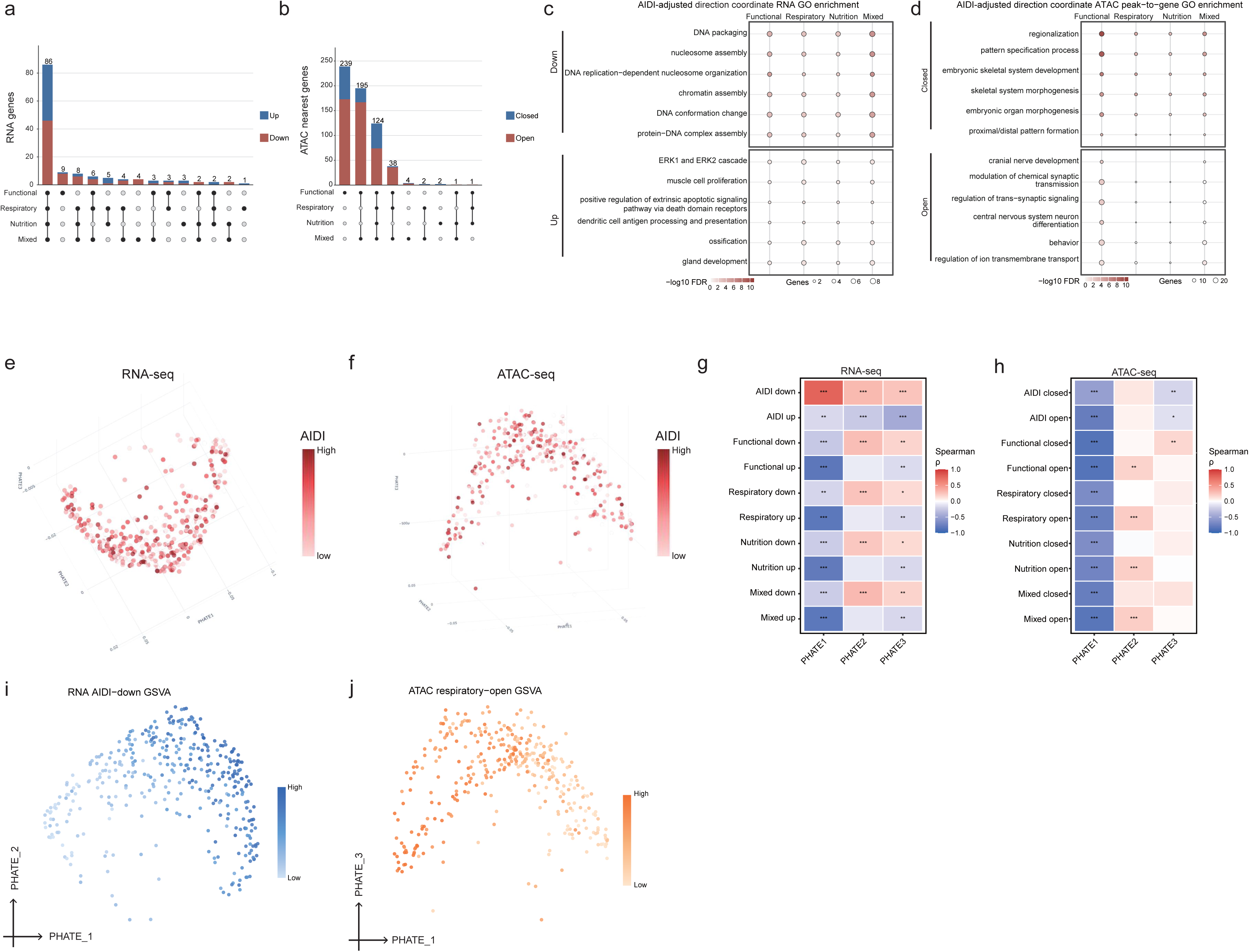
Direction-resolved molecular programs in Answer ALS iPSC motor neurons. a. Exact gene-symbol intersections among four AIDI-adjusted RNA direction sets, split by increasing or decreasing expression. b. Nearest-gene intersections among corresponding ATAC sets, split by higher or lower accessibility. c. Gene Ontology terms for AIDI-adjusted RNA associations with functional, respiratory, nutritional or mixed direction-peak position, split into lower- or higher-expression sets. d. Corresponding nearest-gene terms for AIDI-adjusted ATAC associations, split into closing or opening sets. In b,c dot size shows gene count and colour shows -log10(BH-adjusted P value). Terms were ranked statistically and collapsed for redundancy without biological keyword filtering. e. Three-dimensional PHATE maps of variance-stabilised RNA profiles, coloured by within-sample AIDI rank. f. Three-dimensional PHATE maps of variance-stabilised ATAC profiles, coloured by within-sample AIDI rank. g. Spearman correlations of direction-aligned module GSVA scores with RNA PHATE coordinates. h. Spearman correlations of direction-aligned module GSVA scores with ATAC PHATE coordinates. AIDI modules used direct correlations; Stars show global BH-adjusted significance across the displayed family: * q < 0.05, **q < 0.01 and *** q < 0.001. The heat-map metrics are provided in Supplementary Table S6b. i. Two-dimensional RNA PHATE coloured by the direction-aligned AIDI-down RNA-programme GSVA score. j. Two-dimensional ATAC PHATE coloured by the direction-aligned respiratory-position-associated opening-programme GSVA score.

**Extended Data Fig. 10.**
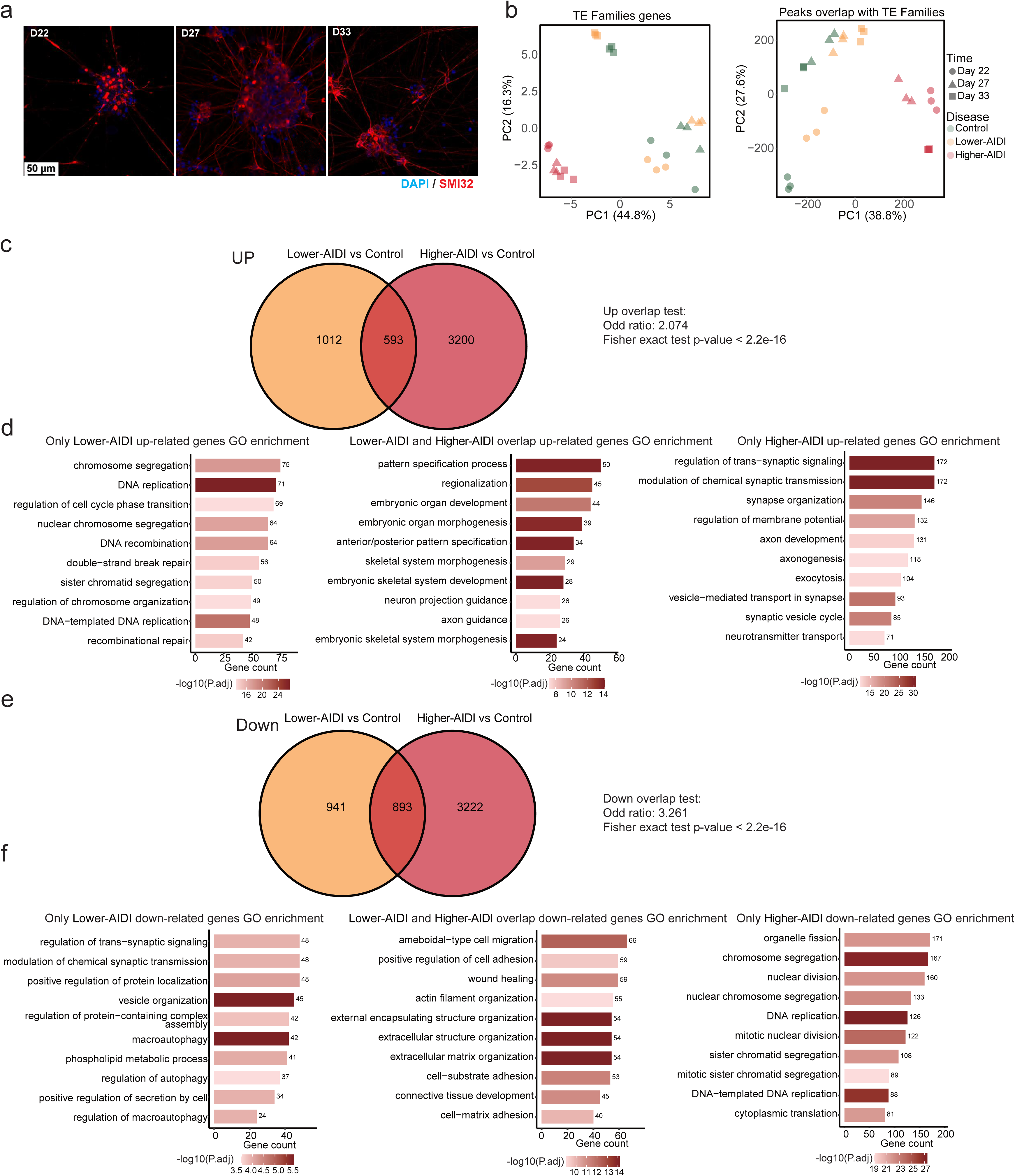
iMN differentiation, TE-restricted variation and persistent expression sets. a. Representative DAPI and SMI32 immunofluorescence images documenting iMN differentiation on days 22, 27 and 33. Scale bar, 50 micrometres. b. Principal-component displays restricted to RNA genes annotated to transposable-element (TE) families (left) or ATAC peaks overlapping TE families (right). Colour shows control, lower-AIDI and higher-AIDI lines; shape shows day. c. Overlap between genes persistently upregulated across all three sampled days in lower-AIDI-versus-control and higher-AIDI-versus-control contrasts. There were 1,012 lower-AIDI-specific, 593 shared and 3,200 higher-AIDI-specific genes (Fisher’s exact test: odds ratio = 2.074, P < 2.2 × 10^−16^). d. Gene Ontology biological-process annotation of the three upregulated partitions. e. Corresponding overlap for persistently downregulated genes. There were 941 lower-AIDI-specific, 893 shared and 3,222 higher-AIDI-specific genes (Fisher’s exact test: odds ratio = 3.261, P < 2.2 × 10^−16^). f. Gene Ontology biological-process annotation of the three downregulated partitions.

**Extended Data Fig. 11.**
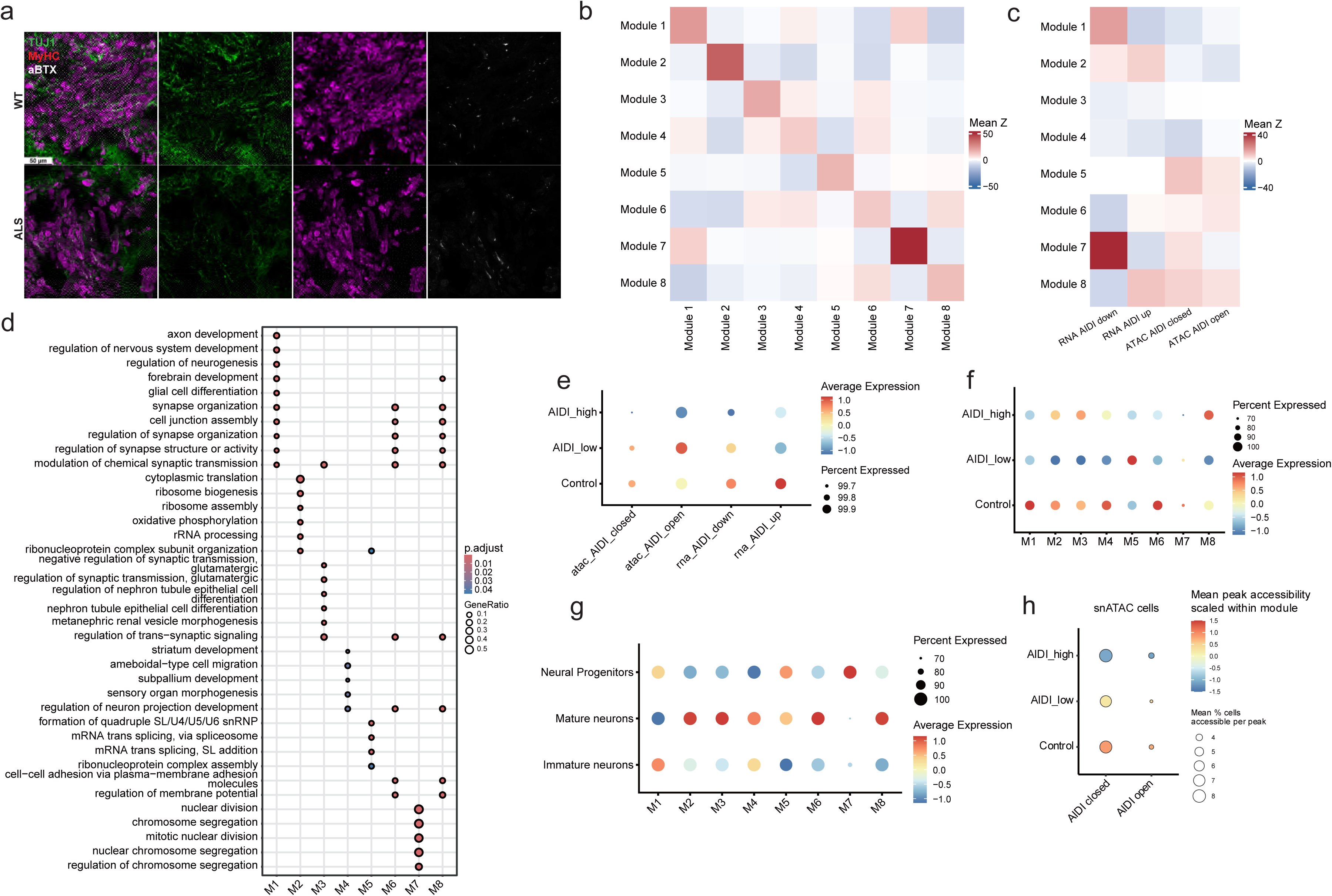
NMO generation and neural-lineage molecular-programme summaries. a. Representative day-50 myosin heavy chain (MyHC) and alpha-bungarotoxin (alpha-BTX) immunofluorescence images documenting generation of control (WT) and ALS neuromuscular organoids (NMOs). Scale bar, 50 micrometres. b. Mean Hotspot local-correlation Z score between each pair of neural-lineage co-expression modules M1-M8, Data are provided in Supplementary Table S7a,b. c. Mean local-correlation Z-score summary between M1-M8 and the transferred AIDI-down RNA, AIDI-up RNA, AIDI-closed ATAC and AIDI-open ATAC programs. The AIDI-down RNA programme aligned most strongly with M7 (mean Z = 25.36) and next with M1 (mean Z = 10.37). Data are provided in Supplementary Table S7a,b. d. Gene Ontology biological-process annotation across M1-M8. M7 was enriched for nuclear-division and chromosome-segregation terms, whereas M1 was enriched for axon development, nervous-system development and synapse organisation. M7 and M1 were positively coupled in 1b (mean Z = 8.61). Dot size shows gene ratio and colour shows BH-adjusted P value. e. Cell-level DotPlot of projected AIDI-associated programme scores in pooled control, lower-AIDI and higher-AIDI neural-lineage cells. f. Cell-level UCell-score DotPlot for M1-M8 by AIDI group. g. Cell-level UCell-score DotPlot for M1-M8 across neural progenitors, mature neurons and immature neurons. h. snATAC-seq DotPlot of accessibility for transferred AIDI-closed and AIDI-open peak sets in neural-lineage cells by AIDI group. Colour shows mean peak accessibility scaled within module; dot size shows the mean percentage of cells accessible per mapped peak.

**Extended Data Fig. 12.**
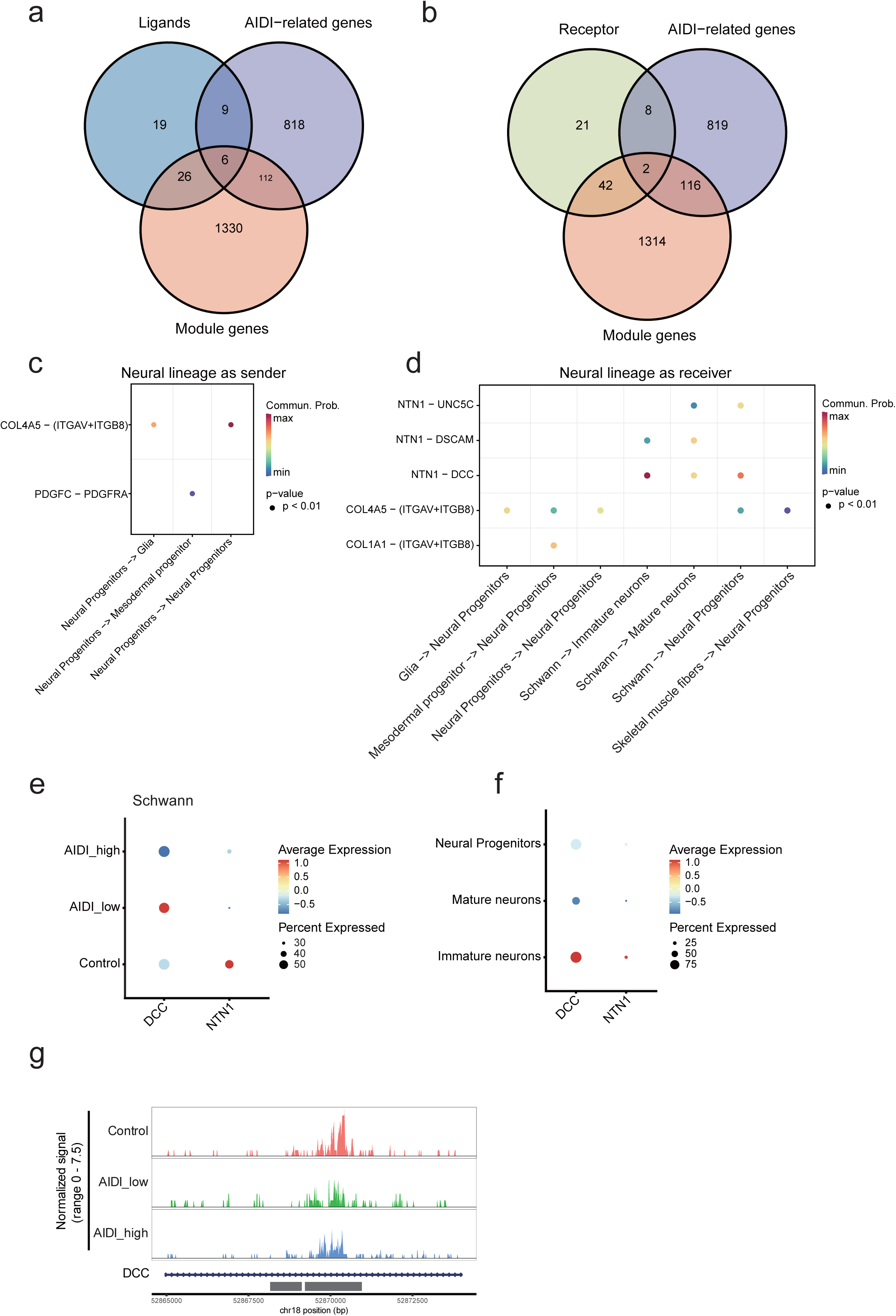
NMO cell-communication and DCC-locus context. a. Intersections of CellChat database ligands with AIDI-associated genes and genes assigned to NMO Hotspot modules. b. Intersections of CellChat database or receptors with AIDI-associated genes and genes assigned to NMO Hotspot modules. c. Selected CellChat-inferred interactions for neural-lineage senders involving ligand genes shared by the AIDI-associated and Hotspot sets. d. Corresponding interactions for neural-lineage receivers, including Schwann-cell-derived Netrin interactions with immature neurons, mature neurons or neural progenitors. e. Cell-level DotPlot of DCC and NTN1 expression in Schwann cells by pooled control, lower-AIDI and higher-AIDI analysis label. f. Cell-level DotPlot of DCC and NTN1 expression across neural progenitors, mature neurons and immature neurons. g. Group-aggregated snATAC-seq normalized accessibility across the DCC locus in immature neurons (hg38, chr18:52,865,000-52,874,000). The prespecified highlighted interval is chr18:52,869,221-52,870,956.

**Extended Data Fig. 13.**
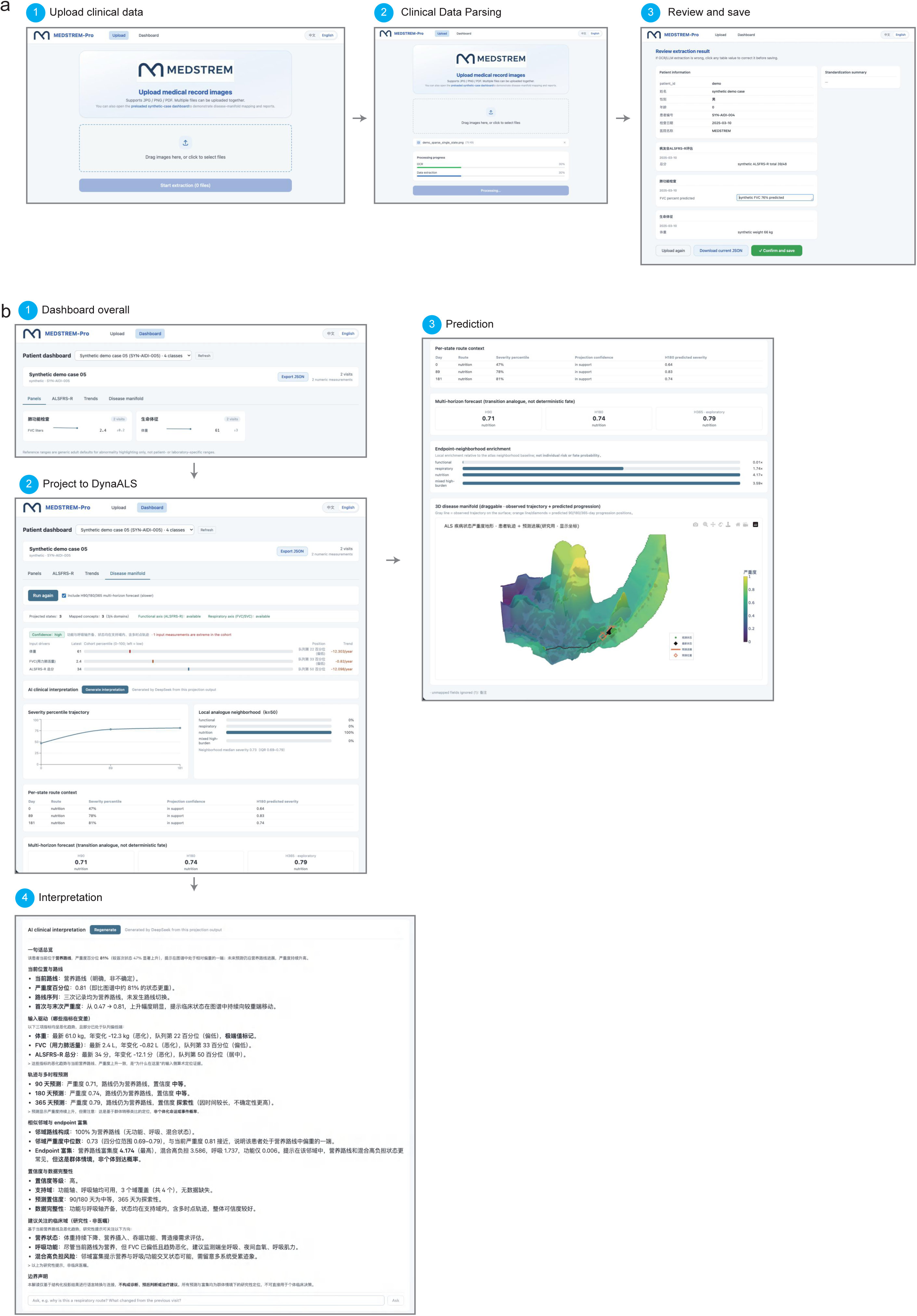
MEDSTREM-App builds and maps an individual longitudinal record. a. Single-user workflow for uploading clinical-record images, monitoring record parsing, reviewing extracted structured fields and confirming or saving the record. b. Application views of a patient dashboard, projection of a structured record onto DynaALS, display of research-use clinical-variable decoder outputs and generation of a text interpretation. The screenshots use demonstration data and show the complete research-use workflow from individual record construction to disease-state mapping.

## Supplementary Table Legends

**Supplementary Table S1. MEDSTREM benchmark performance summary**

Aggregate performance for the four displayed MEDSTREM benchmark stages in Fig. 1c-f. Synthetic image-to-Markdown evaluation used 100 patients, 20 documents per patient and four prespecified image-noise levels. Document-JSON and integrated-record stages retain upstream errors. Real-record image-transcription evaluation used manually deidentified records. Values are arithmetic means; lower character error rate is better and higher key-path accuracy is better.

**Supplementary Table S2. Stage- and document-type-resolved MEDSTREM benchmark**

Benchmark values underlying Extended Data Fig. 3, stratified by synthetic discharge, ECG, EMG, laboratory and vital-sign documents and by prespecified image-noise level. Image transcription, document JSON extraction and longitudinal integration are displayed as a chained workflow; latter-stage values retain upstream errors. The real-record panel reports image-transcription character error rate for deidentified document types.

**Supplementary Table S3. Source-data coverage before analysis-specific eligibility filtering**

a. Distributions of source-file or examination-type coverage per participant for AskHelpU, PRO-ACT and Answer ALS.

b. Calendar-year distribution of AskHelpU medical records.

The most prevalent standardised record or examination types by cohort. Each panel retains its source-specific denominator; the table is descriptive and does not compare cohort quality.

d. clinical variables normalised for use in the current study, with resource-specific fields, units, derivations and applied normalisation.

**Supplementary Table S4. Clinical construct alignment of AIDI**

Same-patient complete-case Spearman correlations of AIDI and three clinical comparator summaries with FVC decline, SVC decline, grip-strength decline and weight loss, underlying Extended Data Fig. 5b. All measures were oriented such that larger values indicate greater deterioration. FVC, SVC and weight loss contributed to AIDI supervision and assess construct alignment; grip strength was excluded from endpoint-specific training and provides a post-fit clinical association. Confidence intervals and paired differences use 5,000 cohort-stratified patient-bootstrap replicates.

**Supplementary Table S5. DynaALS clinical-direction composition and dynamics**

a. Direction-specific state and participant counts together with median and interquartile-range functional, respiratory, nutritional and mixed severity underlying Fig. 3c. The uncertain group is included; participant counts across directions are not mutually exclusive because one participant can contribute states to more than one direction.

b. Nearest-neighbour-matched clinical contrasts between DynaALS directions underlying Fig. 3d. Rows report raw and standardised paired differences, 500 matched-pair bootstrap confidence intervals and BH-adjusted values. These contrasts use patient-time states; patients or comparator states can recur, so they describe direction-specific clinical composition rather than independent patient-level inference.

c. Direction-stratified clinical values across global-burden quintiles underlying Extended Data Fig. 7a. Values are descriptive medians from the full master state set, and colour scaling in the figure is separate within each clinical row.

d. Median and interquartile-range global and domain severity across confidently assigned patient-time states underlying Fig. 3e. Participants can contribute repeated states and can appear in more than one direction.

e. Participant counts and global-severity quartiles stratified by the direction of each patient’s earliest eligible state underlying Fig. 3f. Maximum global severity was calculated from the initial state through available follow-up, inclusive; uncertain initial directions were excluded.

f. observed direction-transition summaries at 90, 180 and 365 days underlying Extended Data Fig. 7b. The table reports row-normalised equal-cohort transition proportions, released numerators and denominators, evaluable participants, uncertainty exclusions and low-count suppression; cells with raw n < 20 remain suppressed.

g. State-level Spearman associations between graph-relative respiratory position and six granular AskHelpU pulmonary-function measurements underlying Fig. 3g. MEF50, PEF, FEV1, MVV, MEF75 and MEF25 are all displayed; the first four reached nominal P < 0.05. State and participant counts are provided for each measurement. P values are nominal and were not adjusted for repeated observations within participants or for multiple pulmonary measurements.

**Supplementary Table S6. Motor-neuron molecular-programs heatmap metrics**

a. Feature counts and false-discovery-rate summaries for unique RNA and ATAC molecular programs underlying Fig. 5c,d. AIDI-adjusted direction-relative programs capture molecular associations beyond global AIDI.

b. Direct or AIDI-adjusted partial Spearman associations between RNA-seq or ATAC-seq PHATE coordinates and programme scores underlying Extended Data Fig. 9g,h. The table preserves the exact association type, adjustment, sample size, rho and global BH-adjusted value displayed in the heatmap.

**Supplementary Table S7. NMO Hotspot modules and frozen molecular programs**

a. Mean local gene-gene correlation Z scores between M1-M8 Hotspot modules underlying Extended Data Fig. 11b.

b. Mean local correlation Z scores between transferred DynaALS-associated RNA/ATAC programs and M1-M8 Hotspot modules underlying Extended Data Fig. 11c.

c. gene membership of Hotspot modules M1-M8 used in Fig. 6h,i and Extended Data Fig. 11b-d.

d. frozen RNA genes and ATAC peaks in the AIDI- and direction-associated programs transferred across the Answer ALS, iMN and NMO analyses. Module correlations describe co-expression structure and do not imply regulatory or causal relationships.

**Supplementary Table S8. Future-state clinical decoding benchmarks**

a. Pooled mean absolute error (MAE) of the predicted future-state decoder and bounded all-visible historical-slope extrapolation underlying Fig. 4e. Each target-horizon comparison used the latest anchor with at least two visible same-target observations spanning at least 30 days, and both methods were evaluated on identical complete cases.

b. Pooled MAE of single-state predicted future-state decoding and an unfitted current-value no-change null underlying Fig. 4f. Only anchor-day information was available, with no earlier individual visit. Intervals use 5,000 patient-cluster bootstrap replicates. The table is restricted to the pooled quantities displayed in Fig. 4e,f.

## Reference

1 Mead, R. J., Shan, N., Reiser, H. J., Marshall, F. & Shaw, P. J. Amyotrophic lateral sclerosis: a neurodegenerative disorder poised for successful therapeutic translation. Nat Rev Drug Discov 22, 185–212 (2023). 10.1038/s41573-022-00612-2

2 Goyal, N. A. et al. Addressing heterogeneity in amyotrophic lateral sclerosis CLINICAL TRIALS. Muscle Nerve 62, 156–166 (2020). 10.1002/mus.26801

3 van Eijk, R. P. A. et al. An old friend who has overstayed their welcome: the ALSFRS-R total score as primary endpoint for ALS clinical trials. Amyotrophic Lateral Sclerosis and Frontotemporal Degeneration 22, 300–307 (2021). 10.1080/21678421.2021.1879865

4 Genge, A. et al. The ALSFRS-R Summit: a global call to action on the use of the ALSFRS-R in ALS clinical trials. Amyotroph Lateral Scler Frontotemporal Degener 25, 382–387 (2024). 10.1080/21678421.2024.2320880

5 Franchignoni, F., Mora, G., Giordano, A., Volanti, P. & Chio, A. Evidence of multidimensionality in the ALSFRS-R Scale: a critical appraisal on its measurement properties using Rasch analysis. J Neurol Neurosurg Psychiatry 84, 1340–1345 (2013). 10.1136/jnnp-2012-304701

6 Roche, J. C. et al. A proposed staging system for amyotrophic lateral sclerosis. Brain 135, 847–852 (2012). 10.1093/brain/awr351

7 Chio, A., Hammond, E. R., Mora, G., Bonito, V. & Filippini, G. Development and evaluation of a clinical staging system for amyotrophic lateral sclerosis. J Neurol Neurosurg Psychiatry 86, 38–44 (2015). 10.1136/jnnp-2013-306589

8 Fang, T. et al. Comparison of the King’s and MiToS staging systems for ALS. Amyotroph Lateral Scler Frontotemporal Degener 18, 227–232 (2017). 10.1080/21678421.2016.1265565

9 Atassi, N. et al. The PRO-ACT database: design, initial analyses, and predictive features. Neurology 83, 1719–1725 (2014). 10.1212/WNL.0000000000000951

10 Baxi, E. G. et al. Answer ALS, a large-scale resource for sporadic and familial ALS combining clinical and multi-omics data from induced pluripotent cell lines. Nature Neuroscience 25, 226–237 (2022). 10.1038/s41593-021-01006-0

11 Ramamoorthy, D. et al. Identifying patterns in amyotrophic lateral sclerosis progression from sparse longitudinal data. Nat Comput Sci 2, 605–616 (2022). 10.1038/s43588-022-00299-w

12 D, M. A., et al. Temporal stratification of amyotrophic lateral sclerosis patients using disease progression patterns. Nat Commun 15, 5717 (2024). 10.1038/s41467-024-49954-y

13 De Mori Bajolin, F., Tavazzi, E., Bianchi, A. M. & Mendez, M. O. Robust end-to-end stratification of amyotrophic lateral sclerosis patients via recurrent variational autoencoder and consensus clustering. J Biomed Inform 180, 105059 (2026). 10.1016/j.jbi.2026.105059

14 Grollemund, V. et al. Development and validation of a 1-year survival prognosis estimation model for Amyotrophic Lateral Sclerosis using manifold learning algorithm UMAP. Sci Rep 10, 13378 (2020). 10.1038/s41598-020-70125-8

15 Zandona, A., Vasta, R., Chio, A. & Di Camillo, B. A Dynamic Bayesian Network model for the simulation of Amyotrophic Lateral Sclerosis progression. BMC Bioinformatics 20, 118 (2019). 10.1186/s12859-019-2692-x

16 Tavazzi, E. et al. Predicting functional impairment trajectories in amyotrophic lateral sclerosis: a probabilistic, multifactorial model of disease progression. J Neurol 269, 3858–3878 (2022). 10.1007/s00415-022-11022-0

17 McFarlane, R. et al. Dynamic modelling of the ALSFRS-R: leveraging population-based scores using neural networks. EBioMedicine 122, 106029 (2025). 10.1016/j.ebiom.2025.106029

18 Tsitkov, S. et al. Disease related changes in ATAC-seq of iPSC-derived motor neuron lines from ALS patients and controls. Nat Commun 15, 3606 (2024). 10.1038/s41467-024-47758-8

19 Hsu, E., Malagaris, I., Kuo, Y. F., Sultana, R. & Roberts, K. Deep learning-based NLP data pipeline for EHR-scanned document information extraction. JAMIA Open 5, ooac045 (2022). 10.1093/jamiaopen/ooac045

20 Luo, M. et al. Automated Extraction of Patient-Centered Outcomes After Breast Cancer Treatment: An Open-Source Large Language Model-Based Toolkit. JCO Clin Cancer Inform 8, e2300258 (2024). 10.1200/CCI.23.00258

21 Wiest, I. C. et al. Privacy-preserving large language models for structured medical information retrieval. NPJ Digit Med 7, 257 (2024). 10.1038/s41746-024-01233-2

22 de Jongh, A. D. et al. Development of a Rasch-Built Amyotrophic Lateral Sclerosis Impairment Multidomain Scale to Measure Disease Progression in ALS. Neurology 101, e602–e612 (2023). 10.1212/WNL.0000000000207483

23 Faghri, F. et al. Identifying and predicting amyotrophic lateral sclerosis clinical subgroups: a population-based machine-learning study. Lancet Digit Health 4, e359–e369 (2022). 10.1016/S2589-7500(21)00274-0

24 Andrews, J. A. et al. Association Between Decline in Slow Vital Capacity and Respiratory Insufficiency, Use of Assisted Ventilation, Tracheostomy, or Death in Patients With Amyotrophic Lateral Sclerosis. JAMA Neurology 75, 58–64 (2018). 10.1001/jamaneurol.2017.3339

25 Marin, B. et al. Alteration of nutritional status at diagnosis is a prognostic factor for survival of amyotrophic lateral sclerosis patients. *Journal of Neurology*, Neurosurgery & Psychiatry 82, 628–634 (2011). 10.1136/jnnp.2010.211474

26 Tavazzi, E. et al. Leveraging process mining for modeling progression trajectories in amyotrophic lateral sclerosis. BMC Med Inform Decis Mak 22, 346 (2023). 10.1186/s12911-023-02113-7

27 Kuffner, R. et al. Crowdsourced analysis of clinical trial data to predict amyotrophic lateral sclerosis progression. Nat Biotechnol 33, 51–57 (2015). 10.1038/nbt.3051

28 Carreiro, A. V. et al. Prognostic models based on patient snapshots and time windows: Predicting disease progression to assisted ventilation in Amyotrophic Lateral Sclerosis. J Biomed Inform 58, 133–144 (2015). 10.1016/j.jbi.2015.09.021

29 Ho, R. et al. Cross-Comparison of Human iPSC Motor Neuron Models of Familial and Sporadic ALS Reveals Early and Convergent Transcriptomic Disease Signatures. Cell Syst 12, 159–175 e159 (2021). 10.1016/j.cels.2020.10.010

30 Ma, G. M. et al. Integrated profiling of iPSC-derived motor neurons carrying C9orf72, FUS, TARDBP, or SOD1 mutations. Stem Cell Reports 20, 102649 (2025). 10.1016/j.stemcr.2025.102649

31 Lee, H. et al. Multi-omic analysis of selectively vulnerable motor neuron subtypes implicates altered lipid metabolism in ALS. Nat Neurosci 24, 1673–1685 (2021). 10.1038/s41593-021-00944-z

32 Liu, E. Y. et al. Loss of Nuclear TDP-43 Is Associated with Decondensation of LINE Retrotransposons. Cell Reports 27, 1409–1421.e1406 (2019). 10.1016/j.celrep.2019.04.003

33 Tam, O. H. et al. Postmortem Cortex Samples Identify Distinct Molecular Subtypes of ALS: Retrotransposon Activation, Oxidative Stress, and Activated Glia. Cell Rep 29, 1164–1177 e1165 (2019). 10.1016/j.celrep.2019.09.066

34 O’Neill, K., et al. ALS molecular subtypes are a combination of cellular and pathological features learned by deep multiomics classifiers. Cell Reports 44 (2025). 10.1016/j.celrep.2025.115402

35 Workman, M. J. et al. Large-scale differentiation of iPSC-derived motor neurons from ALS and control subjects. Neuron 111, 1191–1204 e1195 (2023). 10.1016/j.neuron.2023.01.010

36 Fujimori, K. et al. Modeling sporadic ALS in iPSC-derived motor neurons identifies a potential therapeutic agent. Nat Med 24, 1579–1589 (2018). 10.1038/s41591-018-0140-5

37 Gao, C. et al. Neuromuscular organoids model spinal neuromuscular pathologies in C9orf72 amyotrophic lateral sclerosis. Cell Rep 43, 113892 (2024). 10.1016/j.celrep.2024.113892

38 Ye, L. et al. Sporadic ALS hiPSC-derived motor neurons show axonal defects linked to altered axon guidance pathways. Neurobiology of Disease 206, 106815 (2025). 10.1016/j.nbd.2025.106815

39 Kim, S. H. et al. Axon guidance genes modulate neurotoxicity of ALS-associated UBQLN2. Elife 12 (2023). 10.7554/eLife.84382

40 Hanson, G. et al. Generating real-world data from health records: design of a patient-centric study in multiple sclerosis using a commercial health records platform. JAMIA Open 5 (2022). 10.1093/jamiaopen/ooab110

41 Nottke, A. et al. Validation and clinical discovery demonstration of breast cancer data from a real-world data extraction platform. JAMIA Open 7, ooae041 (2024). 10.1093/jamiaopen/ooae041

42 Wicks, P., Vaughan, T. E., Massagli, M. P. & Heywood, J. Accelerated clinical discovery using self-reported patient data collected online and a patient-matching algorithm. Nat Biotechnol 29, 411–414 (2011). 10.1038/nbt.1837

43 Ma, M. W. et al. Extracting laboratory test information from paper-based reports. BMC Med Inform Decis Mak 23, 251 (2023). 10.1186/s12911-023-02346-6

44 Damani, S. et al. Automating the segmentation, date extraction, and classification of multi-report PDFs in outside medical records using optical character recognition and generative artificial intelligence. JAMIA Open 9, ooag027 (2026). 10.1093/jamiaopen/ooag027

45 Huang, J. et al. A critical assessment of using ChatGPT for extracting structured data from clinical notes. NPJ Digit Med 7, 106 (2024). 10.1038/s41746-024-01079-8

46 Barbiero, P., Vinas Torne, R. & Lio, P. Graph Representation Forecasting of Patient’s Medical Conditions: Toward a Digital Twin. Front Genet 12, 652907 (2021). 10.3389/fgene.2021.652907

47 Huang, Y. & Pepe, M. S. Biomarker evaluation and comparison using the controls as a reference population. Biostatistics 10, 228–244 (2009). 10.1093/biostatistics/kxn029

48 Moon, K. R. et al. Visualizing structure and transitions in high-dimensional biological data. Nat Biotechnol 37, 1482–1492 (2019). 10.1038/s41587-019-0336-3

49 Austin, P. C. Balance diagnostics for comparing the distribution of baseline covariates between treatment groups in propensity-score matched samples. Statistics in Medicine 28, 3083–3107 (2009). 10.1002/sim.3697

50 Hanzelmann, S., Castelo, R. & Guinney, J. GSVA: gene set variation analysis for microarray and RNA-seq data. BMC Bioinformatics 14, 7 (2013). 10.1186/1471-2105-14-7

51 DeTomaso, D. & Yosef, N. Hotspot identifies informative gene modules across modalities of single-cell genomics. Cell Syst 12, 446–456 e449 (2021). 10.1016/j.cels.2021.04.005

52 Jin, S. et al. Inference and analysis of cell-cell communication using CellChat. Nat Commun 12, 1088 (2021). 10.1038/s41467-021-21246-9

